# Multi-dynamic Modelling Reveals Strongly Time-varying Resting fMRI Correlations

**DOI:** 10.1101/2021.06.23.449584

**Authors:** Usama Pervaiz, Diego Vidaurre, Chetan Gohil, Stephen M. Smith, Mark W. Woolrich

## Abstract

The activity of functional brain networks is responsible for the emergence of time-varying cognition and behaviour. Accordingly, time-varying correlations (Functional Connectivity) in resting fMRI have been shown to be predictive of behavioural traits, and psychiatric and neurological conditions. Typically, methods that measure time varying Functional Connectivity (FC), such as sliding windows approaches, do not separately model when changes occur in the mean activity levels from when changes occur in the FC, therefore conflating these two distinct types of modulation. We show that this can bias the estimation of time-varying FC to appear more stable over time than it actually is. Here, we propose an alternative approach that models changes in the mean brain activity and in the FC as being able to occur at different times to each other. We refer to this method as the Multi-dynamic Adversarial Generator Encoder (MAGE) model, which includes a model of the network dynamics that captures long-range time dependencies, and is estimated on fMRI data using principles of Generative Adversarial Networks. We evaluated the approach across several simulation studies and resting fMRI data from the Human Connectome Project (1003 subjects), as well as from UK Biobank (13301 subjects). Importantly, we find that separating fluctuations in the mean activity levels from those in the FC reveals much stronger changes in FC over time, and is a better predictor of individual behavioural variability

**Statement of Significance:**

- MAGE is multi-dynamic in that it models temporal fluctuations in FC independently from fluctuations in the mean of the activity.
- MAGE reveals stronger changes in FC over time than single-dynamic approaches, such as sliding window correlations.
- Multi-dynamic modelling provides an explanation and a solution as to why resting fMRI FC has previously looked so stable.
- MAGE models fMRI data as a set of reoccurring brain states, and importantly, these states do not have to be binary and mutually exclusive (e.g., multiple states can be active at one time-point).
- MAGE estimated time-varying FC is a better predictor of behavioural variability in the resting-state fMRI data than established methods.

## 1. Introduction

Large-scale networks of brain activity can be detected as fluctuations of blood oxygenation levels in functional magnetic resonance imaging (fMRI) [Biswal et al. (1995); Fox et al. (2005)]. These functional networks can be detected in the presence of an experimental paradigm and in spontaneous resting-state activity [De Luca et al. (2006); Tavor et al. (2016); Fox and Raichle (2007)]. Many studies estimate FC by calculating the average correlation across an entire fMRI scanning session data [Rogers et al. (2007); O’Reilly et al. (2012); Smith et al. (2013); Smith (2012); Pervaiz et al. (2020)], and these time-averaged measures of FC have been widely associated with cognitive phenotypes and neuropsychiatric disorders in the literature [Greicius et al. (2004); Anand et al. (2005); Hahn et al. (2011); Hacker et al. (2012); Lynall et al. (2010)].

There is now increasing interest in studying the time-varying nature of FC in fMRI. Significant within-session fluctuations in FC during rest or task fMRI have been demonstrated in a number of studies [Lurie et al. (2020); Vidaurre et al. (2017); Hutchison et al. (2013); Allen et al. (2014); Parr et al. (2018)] and are also well-established in electrophysiological data [Baker et al. (2014); Vidaurre et al. (2018b); De Pasquale et al. (2010, 2012)]. Further, a number of research studies have shown the utility of time-varying FC (TVFC) in identifying many psychiatric and neurological conditions e.g., multiple sclerosis [Van Schependom et al. (2019); d’Ambrosio et al. (2020); Huang et al. (2019)], schizophrenia [Abrol et al. (2017)], Parkinson’s [Cai et al. (2018)], stroke [Chen et al. (2018)], autism spectrum disorder [Guo et al. (2019)], epilepsy [Klugah-Brown et al. (2019)] and major depressive disorder [Zhi et al. (2018)]; and the dynamics of TVFC have been shown to coactivate with spontaneous replay of recently acquired information [Higgins et al. (2021)]. Moreover, inter-subject differences in TVFC have also been associated with a number of behavioural and cognitive traits [Liégeois et al. (2019); Vidaurre et al. (2017)], with some aspects of intelligence only being explained by TVFC [Vidaurre et al. (2021)].

Despite these successes in the estimation and use of TVFC, it is striking how homogeneous FC actually appears to be in fMRI data over time. Indeed, there are a number of studies reporting that there is even insufficient evidence of within-session fluctuations in FC in resting fMRI data [Laumann et al. (2017); Hindriks et al. (2016); Leonardi and Van De Ville (2015)]. While it is possible that close to homogeneous FC in fMRI is a genuine phenomenon, it is also possible that it is instead caused by limitations in the methods used to compute TVFC.

Most previous work is based on the sliding window correlation (SWC) approach [Sakoğlu et al. (2010)]. There are many variants of the SWC method [Allen et al. (2014); Lindquist et al. (2014); Liégeois et al. (2016); Leonardi and Van De Ville (2015)], which are typically limited by using sliding windows of a specified, fixed width. This means that they fail to adapt to the intrinsic time-point by time-point variability of the FC, resulting in changes in FC getting smeared together, thereby making FC look more homogeneous over time. Some of these limitations can be overcome by methods that adapt automatically to time periods of distinct network activity, such as those based on Hidden Markov modelling (HMM) [Vidaurre et al. (2018a); Quinn et al. (2018)]. Even then, a surprising amount of apparent homogeneity in the FC over time persists.

SWC and the HMM have in common that the mean activity and FC estimations are coupled; that is, changes in the mean activity level are by definition accompanied by changes in the FC, when both are modelled. For example, the most common version of the HMM for fMRI characterises each state as a Gaussian distribution with both mean and covariance. Since there is no biological reason for this coupling assumption, in this work we consider the consequences of relaxing that constraint, in fact showing that this change can offer a potential explanation for the homogeneity of TVFC seen in approaches that tie the dynamics of the mean and the FC together.

Our proposed method for uncoupling the dynamics of the mean activity and FC is called the Multi-dynamic Adversarial Generator-Encoder (MAGE). This proposes a multi-dynamic modelling approach that allows the mean activity and FC to fluctuate in time independently from each other. At the same time, MAGE captures long range temporal dynamics of FC and mean activity using recurrent neural networks (in contrast to the Markovian assumption of the HMM) [Gers et al. (2000); Graves and Schmidhuber (2005)]. Finally, our proposed model allows multiple brain states to be simultaneously active (in contrast to the categorical “mutual exclusivity” assumption of the standard HMM). To estimate this model on fMRI data, we developed a novel adversarial generator-encoder network framework inspired by the adversarial regularization used in generative adversarial networks (GANs) [Goodfellow et al. (2014); Lowd and Meek (2005); Makhzani et al. (2015)].

In this paper, we demonstrate the validity of MAGE on several simulation studies and a cognitive task dataset. We also apply the method to resting-state fMRI studies and deduce several outcomes regarding TVFC temporal dynamics, revealing more distinct patterns of FC over time than are obtained with SWC and the HMM, providing an explanation as to why resting fMRI FC has appeared so temporally stable in previous work [Laumann et al. (2017); Hindriks et al. (2016); Leonardi and Van De Ville (2015)]. Finally, we substantiate our findings by demonstrating improved prediction of behavioural and cognitive traits using MAGE-derived TVFC estimates.

## 2. Methods

### 2.1. Multi-dynamic Model

Our proposed approach assumes that fMRI data is generated by a multivariate Normal distribution process with a time-varying mean, *m*_*t*_, and covariance, *C*_*t*_:

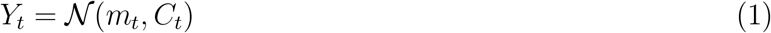

where *Y*_*t*_ is the value of standardised fMRI data, Y (N x T) at time point t, and N is the number of channels (e.g., brain regions).

In order to separately model the time-varying variances and the correlations, we partition the covariance into a NxN diagonal matrix of standard deviations for each brain region, *G*_*t*_, and an NxN correlation matrix, *F*_*t*_:

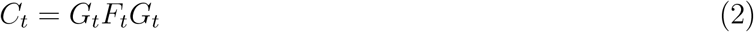

where the correlation matrix, *F*_*t*_, captures instantaneous correlation between brain regions; which we refer to as instantaneous FC.

Clearly, it will not be possible to estimate the instantaneous correlation and means (and variances) at every timepoint without further constraints. As such, we model regularised versions of them by capturing their repeating spatio-temporal structure using an adaptive temporal prior.

Specifically, we assume that the means and correlations (and variances) are generated from their own linear mixture of a limited, underlying set of dynamic modes or states:

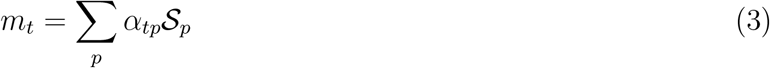

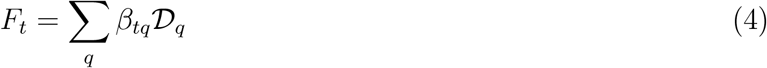

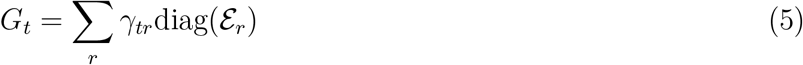

where *α*_*tp*_ is a scalar corresponding to the time course of the *p*^*th*^ “mean activity” state (p=1… P), and 𝒮_*p*_ is its Nx1 spatial map of mean activity; and *β*_*tq*_ is a scalar corresponding to the time course of the *q*^*th*^ “correlation” state (q=1… Q), and 𝒟_*q*_ is its NxN correlation matrix; and *γ*_*tr*_ is a scalar corresponding to the time course of the *r*^*th*^ “variance” state (r=1… R), and ℰ_*r*_ is its Nx1 spatial map of the standard deviations. The 𝒟_*q*_ correlation matrices capture the state-specific correlation between brain regions; which we refer to as the state-specific FC. As we shall describe in detail later, we assume that both the state time courses [*α, β* and *γ*] and the state-specific maps [𝒮_*p*_, 𝒟_*q*_, ℰ_*r*_] are unknown and need inferring from the data.

Note that in Equations (3–5) the correlations and means (and variances) are regularised using their own distinct priors, or dynamics. This is in contrast to most TVFC methods, which are *single-dynamic* approaches in that the dynamics of the means and the correlations (i.e., the FC) are tied together. However, there is no imperative to assume that the dynamics of the means and correlations are the same as each other. As such, the proposed method is designed to explore what happens when we do not tie all of the dynamics together. We refer to this approach as *multi-dynamic*, as illustrated in Figure 1[A].

**Figure 1:**
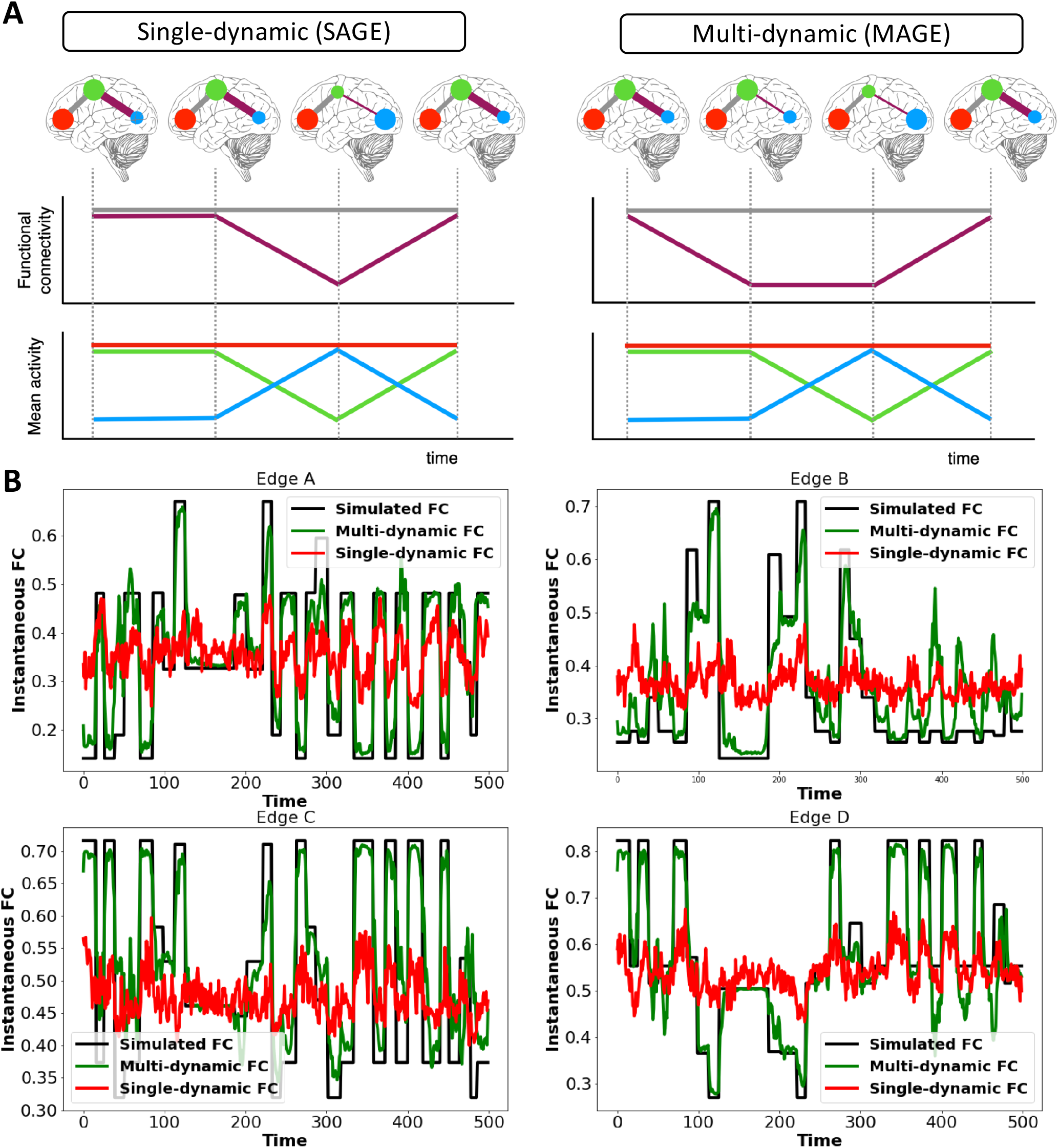
Single-dynamic versus multi-dynamic brain networks. **[A]** Top plot shows example brain network dynamics for the two scenarios; line width indicates functional connectivity (FC) strength, and size of node indicates mean activity amplitude. Middle and bottom plots highlight how the single-dynamic case assumes that the means and the FC change together, whereas in the multi-dynamic case they can change at distinct times. **[B]** Simple simulation illustrating the potential benefits of a multi-dynamic approach. The ground truth is that there are different dynamics underlying the time-varying means and FC. The multi-dynamic approach is able to accurately adjust to fluctuating FC (illustrated for four edges of the FC matrix), whereas the single-dynamic approach is not able to accurately capture these fluctuations, producing time-varying, instantaneous FC that erroneously looks much more homogeneous, as is often observed in time-varying FC estimates in real fMRI data. Lastly, the multi-dynamic approach is also able to adjust to fluctuating mean as shown in Figure S6.

#### 2.1.1. Special case - Sliding Window Correlation

Sliding window correlation (SWC) measures of TVFC capture how the between-brain-region correlation fluctuates over time. For simplicity here (but without loss of generality), we assume non-overlapping sliding windows. This corresponds to Equations (3–5), where *α*_*tp*_, *β*_*tq*_ and *γ*_*tr*_ are set to be the same, and are fixed a priori to correspond to the P=Q=R different positions the sliding window can take as it slides through time (e.g., in the case of square windows, *α* and *β* and *γ* have a value of one inside the sliding window, and a value of zero outside of it). Note that P=Q=R needs to typically be very large to span all the data over time; and so SWC approaches are normally followed by a clustering approach, such as k-means, to find repeating correlation patterns [Allen et al. (2014); Tagliazucchi et al. (2012)]. Typically, information about how the mean fluctuates over time is ignored. Since *α*_*tp*_, *β*_*tq*_ and *γ*_*tr*_ are set to be the same, i.e., the means and correlations (and variances) are modelled using the same dynamics, then SWC corresponds to what we refer to as a *single-dynamics* approach (see Figure 1[A]).

#### 2.1.2. Special case - Hidden Markov Models

Hidden Markov Models (HMM), with multi-variate Normal observation models, capture how both the mean activity levels and the between-brain-region correlations fluctuate over time. This form of the HMM is the same as SWC in that it corresponds to Equations (3–5) where the time courses *α*_*tp*_, *β*_*tq*_ and *γ*_*tr*_ are the same however, *α*_*tp*_, *β*_*tq*_ and *γ*_*tr*_ are now inferred from the data at the same time as 𝒮_*p*_, 𝒟_*q*_ and ℰ_*r*_. This means that the approach can adjust to how the transient network states are coming and going for different durations at different points in time. Note that in addition, the HMM assumes that *α*_*tp*_ = *β*_*tq*_ = *γ*_*tr*_ are binary time courses, defining when states p switches on and off, and that no two states can be active at the same time (i.e., the states are mutually exclusive). As with SWC, since the means and correlations (and variances) are modelled using the same dynamics, then this version of the HMM corresponds to what we refer to as a *single-dynamics* approach (see Figure 1[A]).

### 2.2. What do we mean by functional connectivity?

Typically, correlation-based functional connectivity in fMRI refers to the correlation between two fMRI time series. However, in the model we are proposing, there are multiple entities that could be described as FC, all of which capture some different aspect of correlation between brain regions. First, the model describes the correlation at each time point, i.e., *F*_*t*_ (Equations (1, 2)). We refer to these as the **instantaneous FCs**. Second, a set of state-specific correlations, i.e., *D*_*q*_, are mixed together to regularise the instantaneous FC (see Equation 4). We will refer to these as **state-specific FCs**. Finally, we might also consider correlations in the mean activity time courses, which is a single correlation matrix that corresponds to the correlation of *m*_*t*_ over all time, i.e., Corr(*m*_*t*_). We will refer to this as the **mean activity FC**; this is potentially quite independent of the state-specific FCs.

Our model provides a regularised estimate of the instantaneous FC, and so we can more parsimoniously represent the instantaneous FC using the state-specific FCs. Hence in this work, we focus on using the state-specific FCs as our main TVFC feature of interest, for example when predicting behavioural traits.

### 2.3. Multi-dynamic Adversarial Generator-Encoder (MAGE)

Equations (1–5) describe the modelling framework for the multi-dynamic approach we are proposing. In this section, we will describe the more complete generative model, including the generative process assumed for the dynamics of *α*_*tp*_, *β*_*tq*_ and *γ*_*tr*_, and how we infer on the model. We refer to the overall approach as the *Multi-dynamic Adversarial Generator-Encoder* (MAGE).

In practice, MAGE assumes that the dynamics of the means and variances are tied together, but crucially the correlation is assumed to fluctuate independently. This corresponds to Equations (3–5) with *α*_*tp*_ = *γ*_*tr*_ (P=R), but with *β*_*tq*_ free to be inferred separately^2^. Alongside the state time courses we also infer the state-specific features: 𝒮_*p*_, 𝒟_*q*_ and ℰ_*r*_. Note that while the dynamics of both the means and the variances are always modelled (and are tied together with the same dynamics), for simplicity we will tend to refer to just the dynamics of the means throughout the paper.

In addition, unlike the HMM, MAGE does not assume that state time courses are binary and mutually exclusive. Instead, we assume that they describe time-varying linear mixtures with a sum-to-one constraint (*α*_*tp*_, *β*_*tq*_ and *γ*_*tr*_ separately sums to one at each time point), which we will refer to as “partial volume” state time-courses. This sum-to-one constraint is an identification condition that is imposed to ensure identifiability [Huang (2014)] on the inferred states and makes the post-model temporal analysis interpretable.

In summary, MAGE describes the data as being generated from a finite set of states. These states are linearly mixed together at each time-point to generate a time-varying description of the data’s mean and covariance.

#### 2.3.1. Generative Model

The generative model for the data, *Y*_*t*_, corresponds to Equation 1. Each instance of *Y*_*t*_ is described by a multivariate Normal distribution parameterised by the mean activity at time t, *m*_*t*_, (Equation 3) and the covariance matrix at time t, *C*_*t*_ (Equation 2). We model the state time courses, *α*_*t*_, *γ*_*t*_ and *β*_*t*_, based on their underlying logits:

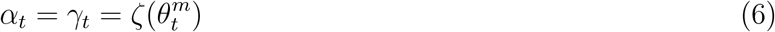

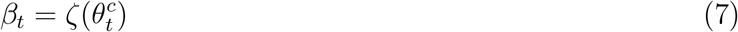

where *ζ* is the softmax function, ensuring that *α* and *β* sum to one (over states) at each time point.

The dynamics in 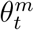 and 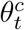 (and implicitly the dynamics of the state time courses) are captured using a unidirectional sequence-to-sequence long short-term memory (LSTM) model, which uses the history of *θ*, to predict the value at the next time point. Specifically, it predicts the prior means of the latent, time-varying parameters 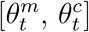.

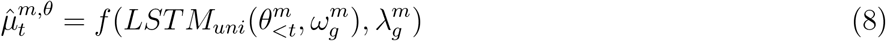

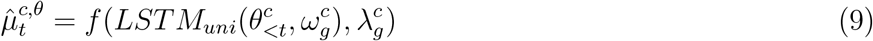

where 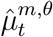 and 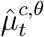 are the prior means of 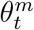 and 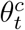 respectively. *f* () is the affine transformation and 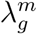 and 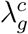 represents the weights of the affine transformation. 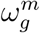 and 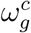 are the generative model LSTM weights. The proposed generative model for the data is illustrated in Figure 2 (and the in-depth architecture is illustrated in Figure S2).

**Figure 2:**
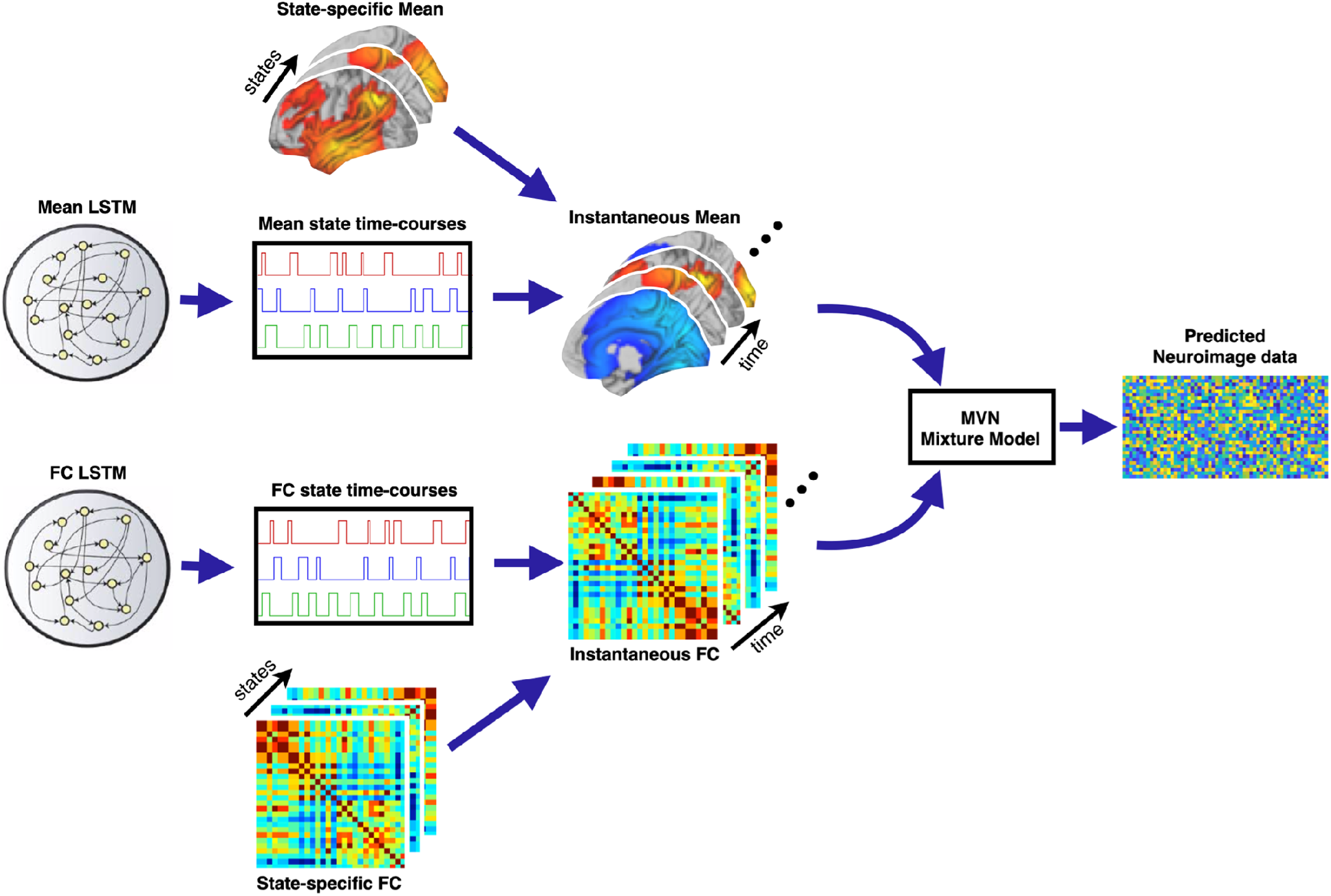
Generative model for MAGE. The proposed model generates data by first generating instantaneous means and the instantaneous correlation matrices (i.e., functional connectivity). MAGE also models instantaneous variances, whose dynamics are tied to be the same as the mean (not shown here). The instantaneous mean is modelled using an underlying set of states, for which the state time courses are generated using a long short-term memory (LSTM) model. The instantaneous correlation is also modelled using an underlying set of states, whose state time courses are generated using a completely different LSTM, making the approach multi-dynamic. The generative model for SAGE is illustrated in FigureS1.

#### 2.3.2. Inference

Given fMRI data, we need to infer on the parameters 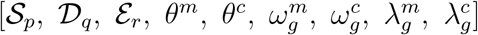 in the generative model. We infer point estimates with uniform priors for the global parameters in the model, i.e., 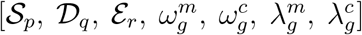; but for the more challenging inference of the latent, time-varying parameters [*θ*^*m*^, *θ*^*c*^], we want to regularise using priors (which here corresponds to the prior means of [*θ*^*m*^, *θ*^*c*^] as stated in Equations (8, 9)). This could be attempted using a number of different approaches. For example, Variational Bayes (VB) methods could be used based on the ideas from Variational autoencoders (VAEs) [Kingma and Welling (2013)], where the generative model acts as a decoder that maps from the latent, time-varying parameters [*θ*^*m*^, *θ*^*c*^] to the fMRI data. The loss function in VB inference is based on optimizing the variational free energy [Friston et al. (2007)] and consists of two terms - the reconstruction likelihood, and the Kullback–Leibler (KL) divergence that regularises the solution by matching the learned posterior distribution to a prior distribution over the latent variables (i.e., Equations (8, 9)).

Here, we take an approach based on an adversarial autoencoder (AAE) [Makhzani et al. (2015)]. Again, the generative model acts as a decoder that maps from the parameters underlying the state time courses, [*θ*^*m*^, *θ*^*c*^], to the fMRI data, and hence the reconstruction likelihood term of the AAE is the same as it would be in a VAE. However, the AAE replaces the regularising KL divergence term of the VAE with an adversarial training criterion [Makhzani et al. (2015); Mescheder et al. (2017)]. This takes the idea of adversarial loss from GANs to achieve the prior regularization - by forcing the posterior means underlying the state time courses, [*θ*^*m*^, *θ*^*c*^], to be close to the specified prior means (i.e., Equations (8, 9)). To do this, we train a discriminator, separately from the training that corresponds to the generative model parameter inference, to be able to distinguish between the prior and posterior versions of the [*θ*^*m*^, *θ*^*c*^]. The trained discriminator is then used to enforce closeness between the posterior and prior (alongside the reconstruction likelihood) when performing the model parameter inference.

##### Encoder Model

The encoder model maps from the fMRI data to the posterior estimates of the latent, time-varying parameters [*θ*^*m*^, *θ*^*c*^]. This can also be referred to as the inference network or inference model and corresponds to the idea of amortized inference in VAEs, whereby the encoder allows an efficient means to “look-up” the posterior estimates given the data [Kingma and Welling (2013)]. The authors who proposed AAEs [Makhzani et al. (2015)] explored the benefits of being fully Bayesian on the latent variables, by assuming that the posterior is a Gaussian distribution (with the mean and variance predicted by the encoder, in a similar manner to the VAE); but after an extensive hyper-parameter search, they did not find any additional advantages, and only reported results using a deterministic version of the posterior. As such, we learned a single nonlinear mapping that is used to map from the data, *Y*_*t*_, to the posterior means at any time point:

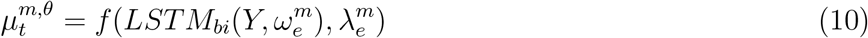

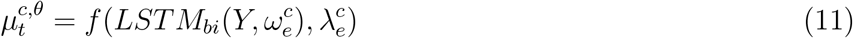

where 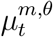 and 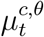 are the posterior means of 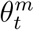 and 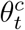 respectively. *LSTM*_*bi*_ is a sequence-to-sequence, bidirectional LSTM model, and 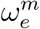 and 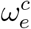 are the encoder LSTM weights. *f* () is the affine transformation and 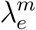 and 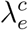 represents the weights of the affine transformation. Note that the approach is still stochastic since we use batching of the data.

##### Discriminator Model

As mentioned above, a discriminator is needed to be able to distinguish between the prior and posterior mean estimates of the state time courses, [*θ*^*m*^, *θ*^*c*^]. The discriminator is also constructed using a sequence-to-sequence, bidirectional LSTM model and it classifies whether samples are from the posterior or prior estimates of [*θ*^*m*^, *θ*^*c*^]:

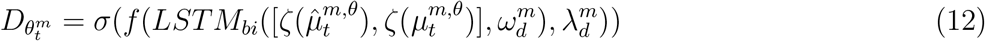

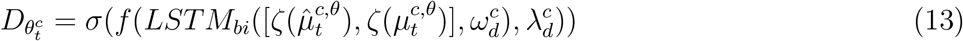

where *σ* is the sigmoid activation to ensure that discriminator output is either 0 or 1, and 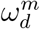 and 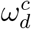 are the discriminator LSTM weights. *f* () is the affine transformation and 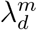 and 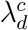 represents the weights of the affine transformation. Note that the discriminator takes in as input the full history of both the prior and posterior mean state time courses into its LSTM. Taken together with the fact that we do not factorise the posteriors of [*θ*^*m*^, *θ*^*c*^] over time, this means that the trained discriminator will help drive the prior and posterior state time courses to have similar characteristics not just over short, but also over long, time scales.

#### 2.3.3. How it Works

We trained this model as a three-player game played between the generator (i.e., the generative model with parameters 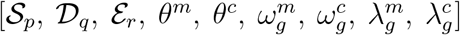), encoder (with parameters 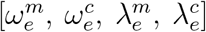), and discriminator (with parameters 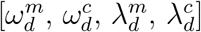). The discriminator model is trained to distinguish prior mean estimates of [*θ*^*m*^, *θ*^*c*^] from their posterior mean estimates. The discriminator is then used to enforce closeness between the posterior and prior of those parameters (while also honouring the reconstruction likelihood) when training the encoder and generator. In this game, when the discriminator successfully identifies the prior from posterior, it is rewarded; whereas both encoder and generator models are penalized. In contrast, when the combined efforts of the encoder and generator models fools the discriminator, the discriminator is penalized. Note, that at the same time the encoder and generator models must also look to maximise the reconstruction likelihood.

This three-player game is played with two main loss functions that are listed below:

1. **Reconstruction (Likelihood) loss:** The first loss corresponds to the likelihood - this tells us how well we are explaining our data according to the current estimate of the generative model, and is given by:

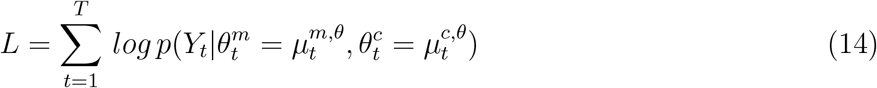
2. **Regularization (Prior) Loss:** The second loss regularises the estimate of the latent, time-varying parameters [*θ*^*m*^, *θ*^*c*^] using an adaptive prior – this penalises when the posterior estimates of [*θ*^*m*^, *θ*^*c*^] deviate from the prior:

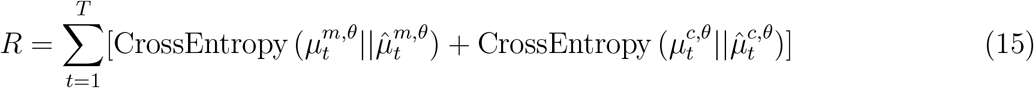 This corresponds to the cross-entropy between the prior and the posterior of the latent, time-varying parameters [*θ*^*m*^, *θ*^*c*^].

The total loss is the summation of these two terms:

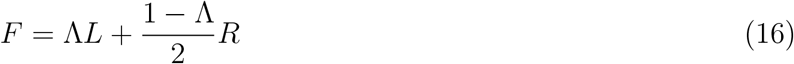

where we attribute a certain weight to each term using Λ. The expression in Equation 16 can be thought of as an approximation of the variational free energy, but where the KL term is replaced by regularisation loss in Equation 15. The complete derivation for the regularisation loss in Equation 15 is given in Section S1.5. We trained the parameters (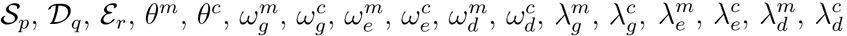) using these losses in two phases - the reconstruction phase and the regularization phase - which are explained below:

- **Reconstruction Phase:** In the reconstruction phase (Equation 14), the generator model (with parameters 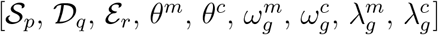) and encoder model (with parameters 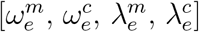) are updated to minimize the reconstruction loss. We kept the discriminator model (with parameters 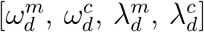) constant during this phase, otherwise the generator and encoder would be trying to hit a moving target and might never converge.
- **Regularization Phase:** In the regularization phase (Equation 15), the discriminator model (with parameters 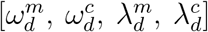) is first updated to distinguish prior and posterior mean estimates of [*θ*^*m*^, *θ*^*c*^]. Then, the parameters of the generator model and encoder model are updated to confuse the discriminator model.

#### 2.3.4. Single-dynamic Adversarial Generator-Encoder (SAGE)

There is variant of the proposed model, SAGE, that assumes the time courses *α*_*tp*_, *γ*_*tr*_ and *β*_*tq*_ are the same. In this regard it is the same as the HMM. However, SAGE maintains the long-range temporal modelling and the partial volume modelling of the network state dynamics that are not part of the standard HMM. Throughout this paper, we will compare MAGE with SAGE to highlight the benefits of the multi-dynamic modelling. SAGE is explained in Section S1.1 and is illustrated in Figure S1.

### 2.4. Dual-Estimation of the group-level MAGE

We performed a process of dual-estimation (analogous to dual-regression following group-ICA) [Beckmann et al. (2009)] to infer the subject-level state versions of the group-level MAGE. This allows us to obtain a subject-specific description of time-varying coorelation and time-varying mean. To achieve this, we used the trained parameters inferred from the group-level MAGE (*α*_*tp*_, *β*_*tq*_, 𝒮_*p*_, 𝒟_*q*_ and ℰ_*r*_). Then we re-estimated the *C*_*t*_ and *m*_*t*_ for each subject by holding constant the *α* and *β* parameters, and only retraining 𝒮_*p*_, 𝒟_*q*_ and ℰ_*r*_.

### 2.5. Hyper-parameters tuning and cross-validation

The hyper-parameters for MAGE were chosen using extensive trial and error by monitoring the validation loss. Some of the hyper-parameters are detailed as follows: 1) The sequence/window length, W, (number of data time-points) was set to 100 and batch-size, B, (number of grouped sequences) was set to 32 - this implies that each batch has a dimension of B*W*N. 2) a small learning rate of 0.0001 was used with large momentum value of 0.9. 3) Nesterov-accelerated Adaptive Moment Estimation (NAdam) optimizer was applied for training the data. 4) the number of units (the dimension of the inner cells in LSTM) for each LSTM layer was set between 8 to 64 (depending on the number of subjects). 5) Λ was optimised depending on the data-set (mostly constrained between 0.85 to 0.95 [Higgins et al. (2016)]). To ensure that the reported MAGE results are reproducible, we performed cross-validations on different non-overlapping sets of subjects. We calculated the similarity of the MAGE estimated state-specific FC correlation and mean activity estimates across these different runs to highlight that the MAGE estimated states are replicable.

## 3. Results

### 3.1. MAGE can infer dynamic linear mixtures of FC states

We start by considering the performance of our proposed method, MAGE, when we set aside the proposed multi-dynamic modelling. This version of MAGE models single-dynamics (where the state time courses for the mean activity and the correlations are the same, i.e., *α*_*tp*_ = *β*_*tq*_) and is referred to as SAGE. SAGE otherwise maintains all of the other features of MAGE, including the long-range temporal modelling and the partial volume modelling of the state dynamics.

We first demonstrate SAGE’s ability to infer state time courses that are dynamic linear mixtures (i.e., partial volume mixtures) with more than one underlying state active at each time point, t. Note that this is in contrast to categorical simulations (i.e., HMM assumptions), where the ground-truth state time-courses, *α* are either 1 or 0 (see Figure S3 for simulations demonstrating M/SAGE’s performance on categorical data). In the partial volume simulations, the ground-truth state time-courses, *α* could be any continuous number from 0 to 1 for any state, but where the sum of *α* for all states, *α*_*jt*_ at any time-point, t, should be 1.

Figure 3[A], shows illustrative examples where the number of simulated states is two and three respectively. Either one, two, or three can be active at any time point. The simulated ground-truth in Figure 3[A] is *single-dynamic* and the simulated state time courses (demonstrated in Figure 3[A]) are common to both the time-varying mean and correlations. We compared SAGE with the HMM to see how they recover the underlying simulated state time courses and the observation model parameters. The HMM failed to infer these state time courses accurately, as HMM is built around the assumption that only one state is active at any time point. On the other hand, SAGE does a good job of inferring these partial volume state time courses. To quantitatively assess the performance of these models, we calculated the Riemannian distance between the simulated covariances and the inferred covariances for each time point. The lower the Riemannian distance between the inferred and simulated covariance, the better the performance of the model. We can see that SAGE can accurately infer the underlying time-varying covariance, but this is not the case for the HMM.

**Figure 3:**
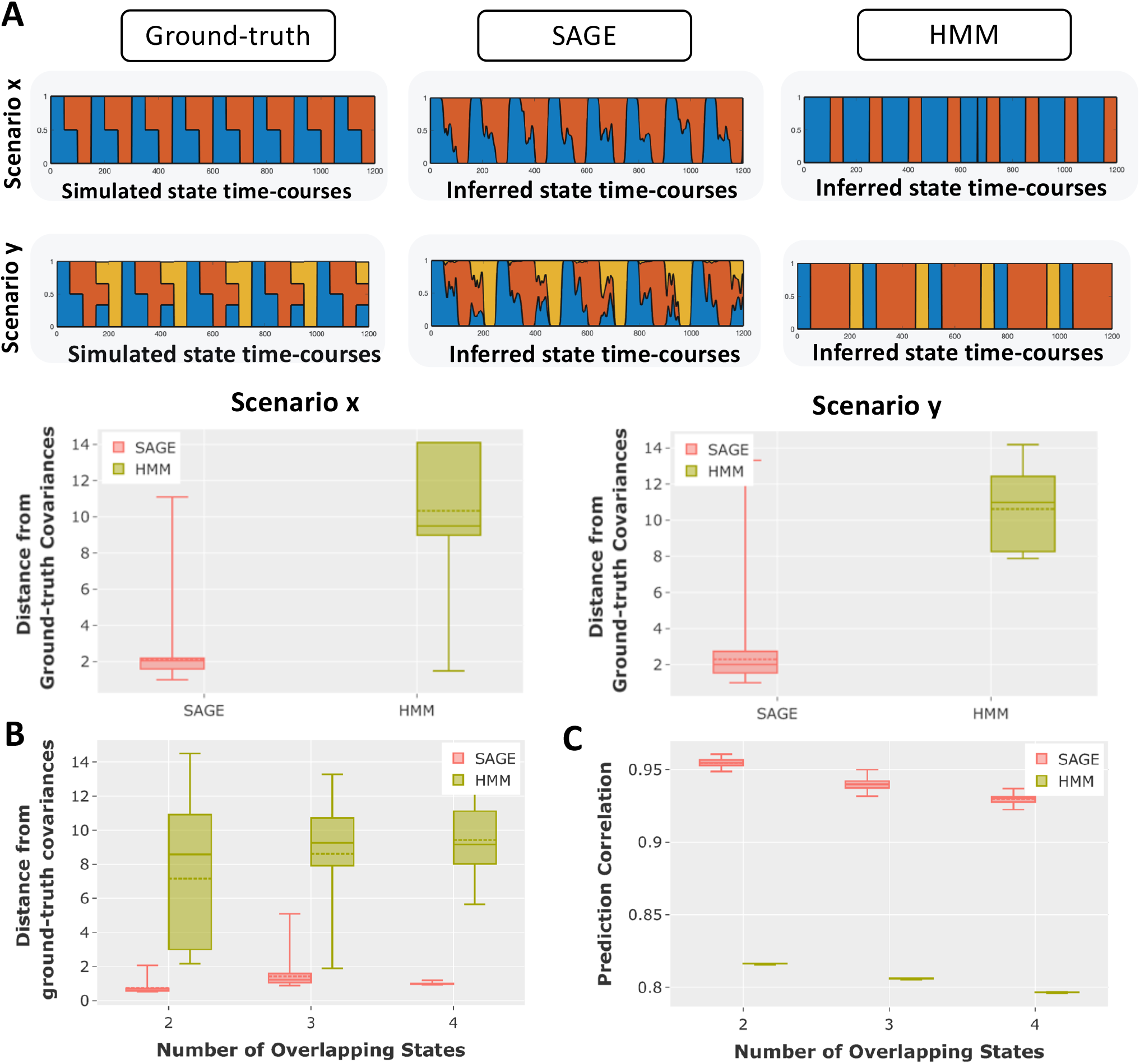
Comparison of the performance of the single-dynamic approach SAGE versus HMM, using simulations of dynamic linear mixtures of FC states (i.e., partial volume) and single-dynamics dynamics (i.e., where the state dynamics of the means and correlations are tied together). **[A]** Two illustrative partial volume examples, comparing SAGE performance with HMM. In scenario x, two states can be simultaneously active, whereas, in scenario y, up to three states can be simultaneously active. Note that these state-time courses were common to both the time-varying mean and correlations (i.e., instantaneous FC). The bottom row of **[A]** shows the Riemannian distance between the inferred and simulated covariances for both HMM and SAGE. **[B-C]** Comparison of the performance of SAGE with HMM on a number of partial volume simulations. Each box plot shows the distribution of prediction performance results over 50 different simulations. Riemannian distance between inferred and simulated covariances was used as a measure of success in **[B]**, and prediction correlation between the inferred and simulated state time-courses in **[C]**.

Figure 3[B,C] shows the performance of the approaches across many partial volume scenarios. We varied the number of overlapping states (either 2, 3, or 4), and then compared the performance of SAGE against the HMM. For each overlapping state configuration, we generated 50 different simulations in which we varied the state time courses (% of overlap), and the observation model itself. We illustrated the Riemannian distance between the inferred and simulated covariances. We also showed the prediction correlation (not accuracy, as state time courses, are no longer categorical) between the inferred and the simulated state time courses.

Lastly, we demonstrate the performance of MAGE for scenarios where the ground-truth is such that FC is static (i.e., no fluctuations in edges of the FC matrix over time). We show that MAGE does not infer time-varying correlations when we infer more than one state for such simulations (see Figure S4). This highlights that MAGE only infers time-varying FC when the underlying FC is time-varying.

### 3.2. Failing to model distinct mean activity dynamics can result in homogeneous estimation of FC over time

Commonly-used SWC and HMM approaches are what we refer to as single-dynamic models, as they tend to tie the dynamics of the correlations (i.e., the FC) to be the same as the dynamics of the means of the brain activity. However, if the underlying truth is that the dynamics of the means and correlations are actually distinct from each other, then using single-dynamic approaches to infer TVFC can have negative consequences. This is because forcing *α*_*tp*_ to be the same as *β*_*tq*_, when the ground truth is that they are not the same, causes the dynamics used for estimating the state-specific FC to be incorrect. This then leads to a smearing of the FC estimation over time, making it look more homogeneous than it really is.

We illustrate the mechanism of the multi-dynamic approach using a simple scenario in Figure 1[A]. Furthermore, in Figure 1[B], we demonstrate the potential benefits of the multi-dynamic approach on a simulation ^3^ in which the ground truth is such that the correlation dynamics and the mean activity dynamics are not tied together. Specifically, Figure 1[B] shows the instantaneous FC estimated using SAGE or MAGE when we simulate using equations (1–5) with *α*_*tp*_ not equal to *β*_*tq*_.

When we infer TVFC using SAGE, the single-dynamic version of MAGE (with *α*_*tp*_ set to be the same as *β*_*tq*_), there are substantially reduced fluctuations in the inferred instantaneous FC over time (i.e., FC is mapped as being near-homogeneous/static) as is often observed in TVFC estimates in real fMRI data. In contrast, the multi-dynamic version of MAGE (with *α*_*tp*_ free to be not equal to *β*_*tq*_) is better able to capture the true fluctuations in instantaneous FC over time.

### 3.3. MAGE can infer dynamics in the FC that are distinct from dynamics in the means of brain activity

We next sought to see if MAGE could identify multi-dynamics in a range of scenarios. For this, we simulated distinct dynamics (i.e., multi-dynamics) for the mean and correlations (i.e., the FC) by simulating two different state time courses, *α*_*tp*_ and *β*_*tq*_, using two different hidden semi-Markov model chains, whose state lifetime values were sampled from two different gamma distributions such that *α*_*tp*_ was not equal to *β*_*tq*_. These state time courses were then used in equations (1–5) to simulate the data.

We compared the performance of the multi-dynamic approach (MAGE) with the single-dynamic approach (SAGE) in Figure 4[A,B]. As illustrated in Figure 4[A], the lifetimes of both the mean and correlation states in the simulation were sampled from the gamma distribution. However, we fixed the distribution of lifetimes of the correlation state time courses by setting the gamma distribution parameter to k=30, whereas, for the mean state time courses, we varied k from 5 to 80, so that the mean state time courses spanned a variety of timescales in different scenarios. In Figure 4[B], it is the other way around, where we fixed the lifetime distribution of the mean state time courses (k =30), but varied the lifetime distribution for the correlation state time-courses (again from k = 5 to 80). In Figure 4[A,B], we have plotted the prediction accuracy for the state time courses of correlation and mean. MAGE does a better job in predicting the underlying simulated labels for state time-courses as compared to SAGE. Note that we did not draw any comparison with the HMM here, as the HMM is also a single-dynamic approach, and the HMM is not able to infer the mean and correlation in such a multi-dynamic fashion (and hence the results would be expected to be closest to SAGE).

**Figure 4:**
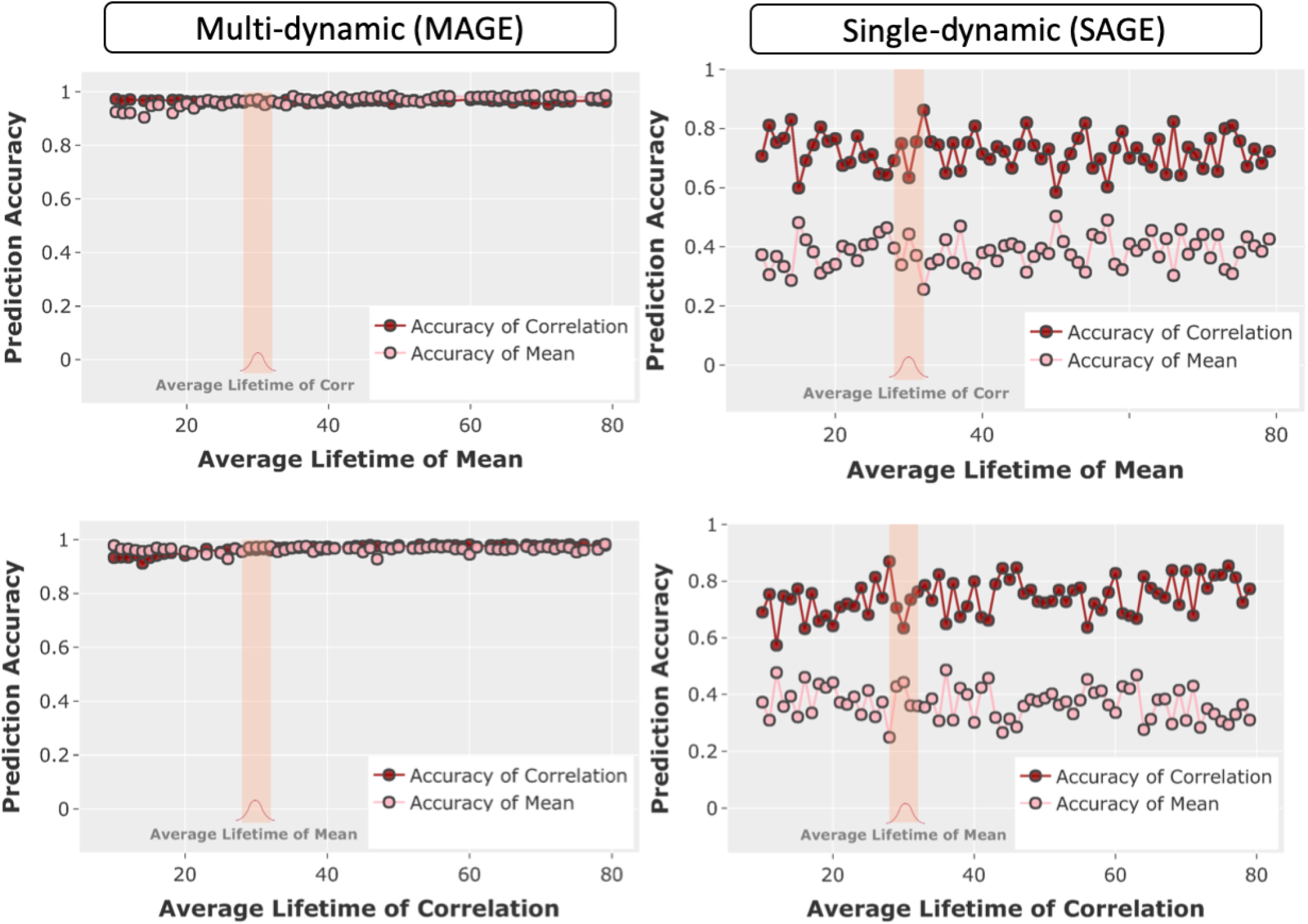
Comparison of the performance of the multi-dynamic approach (MAGE) and the single-dynamics approach (SAGE) on multi-dynamic simulations where multi-dynamics are the ground truth (i.e., where the state dynamics of the means and correlations are free to fluctuate independently). **[A**,**B]** The prediction performance of multi-dynamic and single-dynamic approaches are compared across a range of simulations. The x-axis shows the lifetime parameter (sampled from a gamma distribution) for the mean and correlation state time-courses, whereas the y-axis shows the prediction accuracy of the inferred state time-courses. Specifically, in **[A]**, we fixed the lifetime distribution of the correlation state time-courses (k = 30, as shown by the shaded bar) but varied the lifetime distribution for the mean state time-courses (from k = 5 to 80), and it is the other way around in **[B]**.

### 3.4. MAGE learns network state dynamics that show appropriate task dependencies

In order to help validate the different dynamics that MAGE is inferring, we applied it to task fMRI data. The idea is to train MAGE with no knowledge of the task timings (i.e., unsupervised), allowing us to see if the state dynamics being inferred by MAGE are meaningful in the sense that they exhibit dynamics that link to the task. Furthermore, we wanted to compare how the FC varies across the task conditions using the MAGE approach (and judge whether the inferred spatial maps are meaningful).

Task fMRI data used was collected as part of a previous study from 15 healthy volunteers (7 females, 8 males, age = 27.3±4.4yr, all right-handed) without any previous neurological disorders [Kieliba et al. (2019); Duff et al. (2018)]. Participants performed four different tasks: rest, motor only, visual only, and simultaneous (but independent) visual and motor tasks, and each task was repeated four different times. The motor task involved continuous finger-tapping against the thumb using the right hand. The vision task consisted of videos of colourful abstract shapes in motion. During the combined motor-visual task, participants performed the finger-tapping task while simultaneously watching the videos. We applied a framework used for identifying large-scale probabilistic functional modes (PROFUMO) [Harrison et al. (2015, 2020)], to identify parcels that are allowed to be correlated with each other in space and/or time, and which explicitly models between-subject variability in the parcels. We applied PROFUMO on this data with a dimensionality of 30, which outputs 30 parcel time-courses. These then correspond to the N=30 channels of data that are fed into MAGE.

#### Temporal Dynamics

In the main results shown in Figure 5, we modelled four states using MAGE, so that the number of states is matched to the number of task conditions. In Figure 5[A], we have plotted the task-evoked occupancy, which is the average of the MAGE inferred state time courses across the four repeated runs (as each experimental condition is repeated four times). The task-evoked occupancy shows clearly that the inferred states switch at the same time as the experimental task condition changes. It is worth noting from Figure 5[A] that the estimated FC-driven state-time courses, *β*_*tq*_, performed better than the mean activity/variance-driven time-courses, *α*_*tp*_. Moreover, in Figure S7, we have illustrated the results of task decoding when we modelled more than 4 states. In Figure S7[A], we have modelled 8 states and in Figure S7[B], we have modelled 16 states, and we plotted the relative fractional occupancy (time spent in each state) distribution for each of the modelled states. It is evident that even with the increased number of states, we can still see dynamics that are task dependent and can identify task-specific states (as highlighted by the paired t-test values on the respective states).

**Figure 5:**
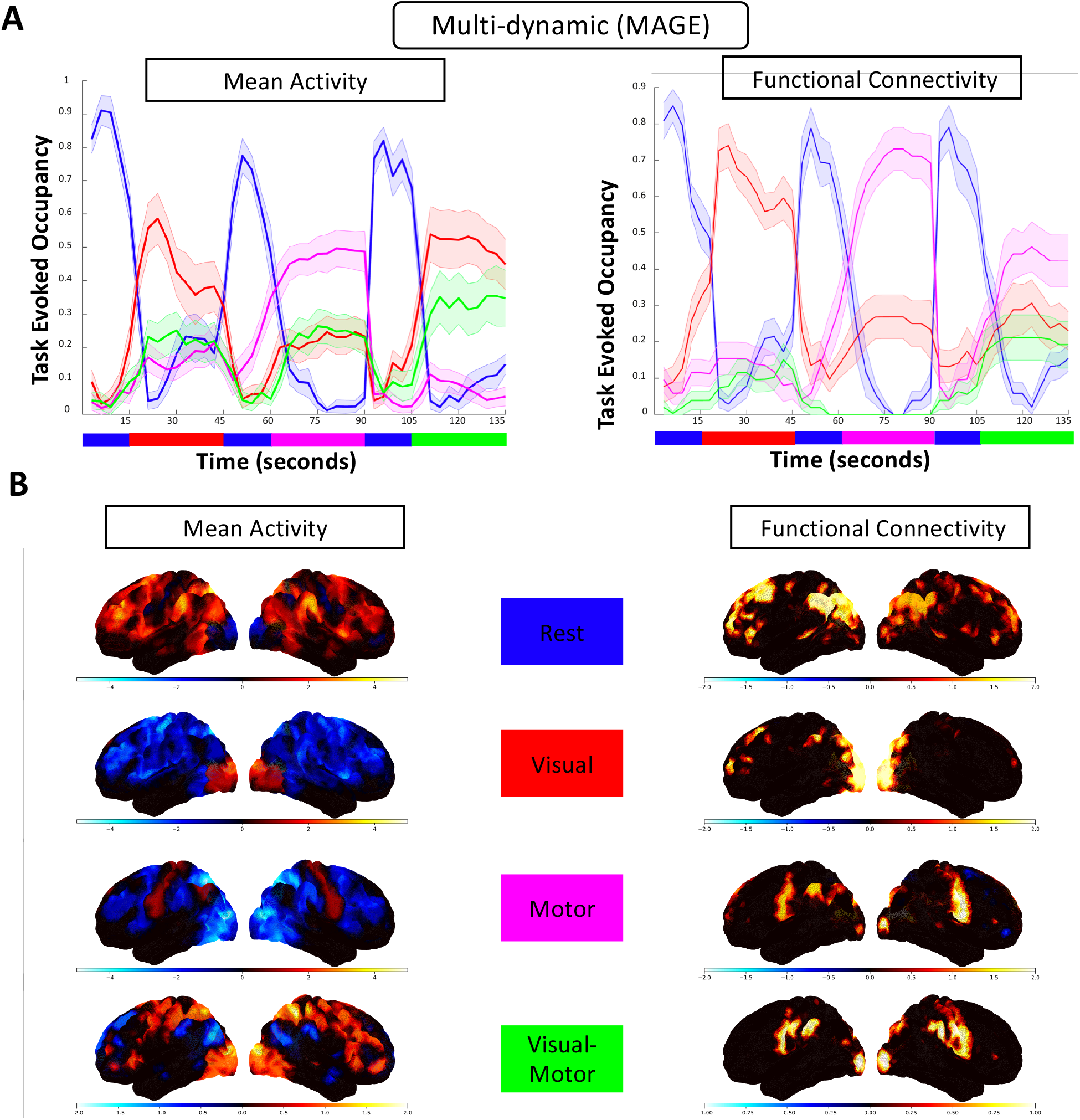
**[task fMRI data]** Validation of the performance of MAGE on a on a real task fMRI data-set. **[A]** There are four different tasks performed by the participants (rest, visual, motor, and visual-motor). Task-evoked occupancy plots of the state time courses from MAGE (learnt with no knowledge of the task timings) across the four tasks, where the x-axis represents the number of time points and the y-axis corresponds to the average activation value for that state. Specifically, the task-evoked plot curves show the mean and standard deviation across four repeated runs of each experimental condition for all 15 subjects. **[B]** The state-specific spatial maps for the functional connectivity (FC) and mean activity, which are labelled according to the task conditions that their state time courses most correspond to. The state-specific spatial maps for variance are not illustrated as there is no between-state variability in the inferred variance states. For the mean activity maps, the average amplitude of each node/channel is shown. The negative mean activity (e.g., deactivation of the visual network during the motor task) refers to decreases in signal change compared to the baseline. For the FC maps, rank-one decompositions (first Eigenvector) of the estimated state-specific FC correlation matrices are shown. The magnitude of the values of the state-specific FC spatial maps (colorbars) correspond to the degree of connectedness of each region with the rest of brain.

#### Spatial Dynamics

In Figure 5[B], we show the state-specific FC spatial maps (corresponding to the the first eigenvector of the state-specific correlation matrices) and mean activity spatial maps for each of the four inferred states. The variance spatial maps are not illustrated as there was no between-state variability in the inferred variance states (lack of time-varying content in variance). The maps are labelled according to the task conditions that their state time courses most correspond to. The state-specific FC maps for MAGE identifies appropriate active regions during each of the tasks. It can be seen during the motor task that the somatomotor area of the brain is active, and during the visual task that the visual cortex is active. During the rest task, the Default Mode Network (DMN) is active, and during the visual-motor task, both the motor and visual cortex are active. The mean activity level maps appear fairly similar and look appropriate with regards to the relevant task condition. During the visual task, the visual cortex of the brain is active, and during the motor task, the motor cortex is active for both approaches. During the rest task, DMN is active, and during the visual-motor task, only the visual cortex is active for the MAGE approach.

### 3.5. Multi-dynamic approach reveals stronger changes in FC over time in resting fMRI

The simulations shown in Figure 1 suggest that multi-dynamic approaches may be better able to identify TVFC than single-dynamic approaches, which may erroneously infer FC as being too homogeneous over time. We looked to test this idea by applying the single-dynamic (SAGE) and multi-dynamic (MAGE) approaches to resting-state fMRI data. In particular, we can analyse how the temporal dynamics of mean activity levels and FC differ (i.e., *α*_*tp*_, *β*_*tq*_) when modelled independently and see if this has any effect on the inferred state-specific FC maps. Lastly, we compared MAGE results with the HMM and SWC approaches, to further substantiate our findings.

We used rfMRI data from 13301 subjects from the UK Biobank (UKB) [Miller et al. (2016)] and 1003 subjects from the Human Connectome Project (HCP) [Van Essen et al. (2013)]. The pre-processing pipelines for UKB and HCP main steps can be summarized as (1) motion correction, (2) removal of structural artifacts with ICA (Independent Component Analysis and FMRIB’s ICA-based X-noisefier (FIX), and (3) brain parcellation using Group-ICA. The length of the scanning session for UKB is 6 minutes, and for HCP is 1 hour (4 sessions of 15 mins each). For UKB, we have 490 time-points and for HCP, we used data from all 4 sessions which gave us 4800 time points in total (1200 time points in each session). Lastly, we applied a popular data-driven parcellation scheme called spatial Group Independent Component Analysis (groupICA) to our data to identify the parcels/channels (N) that form the data to be input into MAGE.

#### Temporal and Spatial Dynamics

We first compare the single-dynamic (SAGE) and multi-dynamic (MAGE) approaches using the HCP data, where the only difference between the two methods is whether or not we model independent state dynamics for the state means and FC (i.e., correlations). In Figure 6[A], we show the state-specific mean activity and FC correlation maps for SAGE and MAGE approaches for the HCP data. In total, we modelled P=Q=R=12 states, but we have only displayed the observation model for four of the 12 states for ease of visualisation (the results for all 12 states are illustrated in Figure S9, Figure S10, Figure S11 and Figure S12). As described earlier, MAGE also models time-varying variances, whose dynamics are tied to be the same as the mean. However, we have not displayed the state-specific spatial maps for variance as MAGE infers that there was little between-state variability in the state-specific variances (i.e., the variances remained static across states).

**Figure 6:**
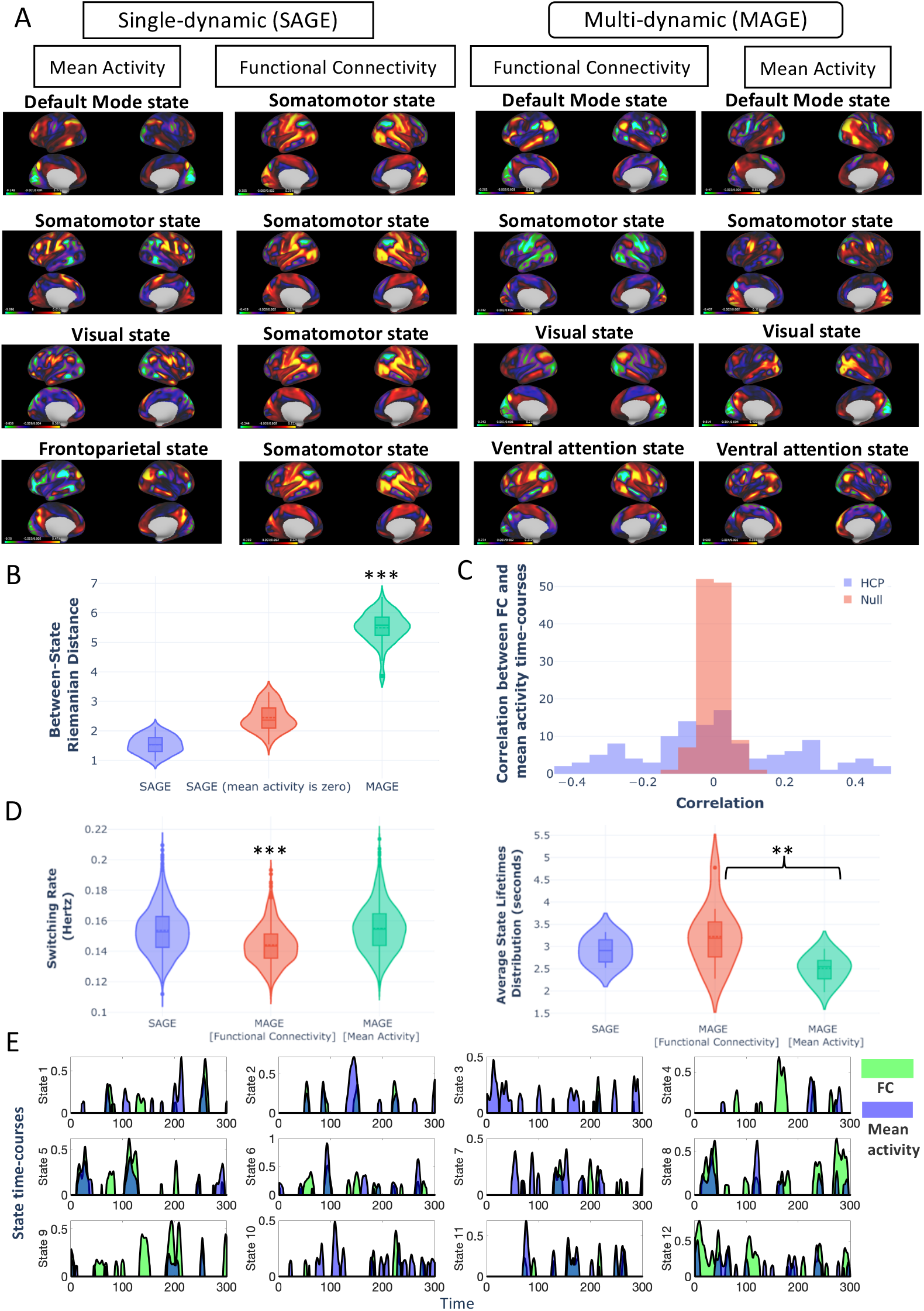
**[HCP data] [A-B]** The multi-dynamic (MAGE) approach reveals stronger changes in state-specific FC maps compared to the single-dynamic (SAGE) approach. Using the HCP data, it is evident that qualitatively and quantitatively, MAGE-derived FC shows stronger changes over time. **[A]** State-specific mean activity and FC correlation maps for four of the modelled states for both approaches are shown. For the state-specific mean activity maps, the average amplitude of each node/channel is shown and for the state-specific FC maps, rank-one decomposition (first Eigenvector) of the estimated FC correlation maps are shown. **[B]** The pair-wise Riemannian distance of the state-specific FC correlation matrices between 12 FC states is shown for both MAGE and SAGE (with and without modelling the mean activity, see [Vidaurre et al. (2021)] as another example of modelling only the correlation, i.e., *m*_*t*_ = 0 in Equation 1). **[C]** Correlations between the mean-derived state time-courses and the FC-derived state time-courses plotted as histogram over all pairs of states (and equivalent null scenario). The non-zero correlations indicate that the mean activity and FC have some shared dynamics but also have unique temporal dynamics. **[D]** Summary measures of the state dynamics for both approaches. It can be seen that FC state dynamics show the lowest switching rate (in Hertz) and the highest lifetimes values (in seconds) as compared to other approaches. **(E)** Example segment of a state time course of a randomly chosen subject is shown to qualitatively highlight the extent of similarity (and dissimilarity) between the mean activity and the FC state time courses. These also show how states in MAGE can overlap in time.

Figure 6[A] shows that the state-specific mean activity maps vary reasonably well across states using both SAGE and MAGE. However, with SAGE, it can be seen that the state-specific FC maps (the rank-one decomposition of the estimated state-specific correlation maps are shown to make these 2D FC maps easier to view) of all the states are predominantly similar, always resembling a somatomotor state. We also attempted another modelling option with SAGE in which we explicitly forced the mean activity to be zero^4^ (i.e., *m*_*t*_ = 0 in Equation 1). This approach also resulted in homogeneous state-specific FCs as shown in Figure S13. In contrast, for the MAGE approach, we can see that the state-specific FC maps correspond to distinct brain networks, with the state-specific FCs looking more distinct from each other than in the SAGE results. To illustrate this quantitatively, we show the pairwise between-state Riemannian distance in Figure 6[B]. Specifically, we calculated these pairwise distances over the actual 2D correlation matrices (i.e., FCs). It is clear that these between-state pairwise distances are higher for the state-specific FC maps when inferred using the multi-dynamic MAGE approach, revealing stronger changes in FC over time.

In Figure 6[D], we compare summary measures of the state dynamics. Since MAGE models a linear mixture (i.e., a partial volume) at each time point, it is not straightforward to compute summary statistics normally used on binary state time courses. In order to allow us to do this we threshold the continuous partial volume state time courses (*α*_*tp*_ and *β*_*tq*_) at 0.3. Firstly, we show the state switching rate for both models, which is a measure of the frequency of state fluctuation for each subject, and it can be seen that FC-driven state time-courses, *β*_*tq*_, have the lowest switching rate. Then, we show the state lifetimes for both models, which correspond to the number of seconds per state visit. We can see that the FC-driven state time-courses have the highest values for state lifetimes (2 to 5 seconds) as compare to lifetimes of mean state time-courses (2 to 3 seconds). In Figure 6[E], we show an example segment of the state time course (of a randomly chosen subject) showing the nature of the mixing of the inferred 12 states and the degree of overlap between the mean activity and the FC state time-courses. Furthermore, in Figure 6[C], we quantitatively compare the extent of similarity between the mean activity and the FC state time courses. We plot the mean activity and the FC state time courses correlation matrices as histograms for the null and the HCP dataset. There is a significant non-zero correlation as compared to the equivalent null scenario, but there is clearly still a distinct difference between them (and hence the difference in the performance of SAGE vs MAGE). In short, the means and FCs have some shared dynamics but also some unique/independent dynamics.

It has been previously proposed that the observed estimates of TVFC must be compared against “static” null hypotheses to establish claims of non-stationary [Lurie et al. (2020)]. Considering that, we generated null data using autoregressive randomization (ARR) following [Liegeois et al. (2017)]. Specifically, a 10-th order Gaussian autoregressive (AR) model was applied to generate null data. For each participant and each pair of brain regions, we compared the observed between-timepoint variances of the instantaneous FCs inferred by MAGE, which can be calculated from the FC state dynamics and state-specific FCs, from the real HCP data against the null data. This analysis resulted in one p-value for each edge of the FC matrix. On average, across HCP participants (compared against null data), 95% of edges were significant (p *≤* 0.001). Furthermore, in Figure S5, we show the histogram of null variance values compared with the real HCP data variance values (this highlights that between-brain region variability over time is significantly higher in the real HCP data as compared to null data).

We find that we get similar results to the HCP dataset when we use MAGE on the resting fMRI data from the UKB dataset. Here, we also compare the performance of MAGE with other established methods (HMM and SWC) on the UKB data-set. Figure 7 shows the state-specific FC maps from MAGE, HMM and SWC, with the HMM and SWC approaches showing similar behaviour to that of SAGE in Figure 6. It is also clear that there are smaller between-state differences in FC across time for both the HMM and SWC approaches compared to MAGE (as reflected by the between-state Riemannian distance plots in Figure 7[D]), consistent again with the idea that MAGE reveals stronger changes in FC over time due to its ability to model multi-dynamics.

**Figure 7:**
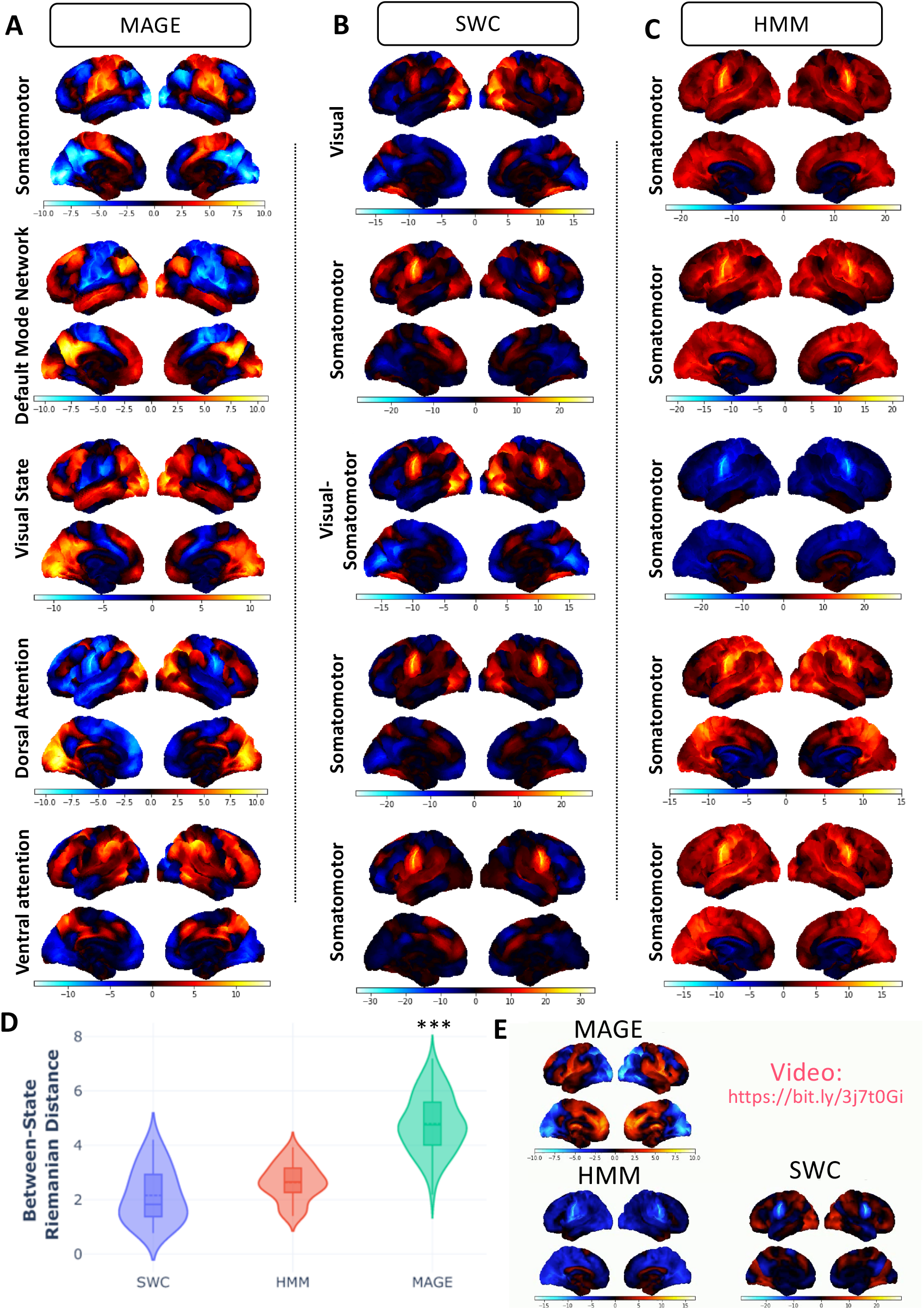
**[UKB data]** Demonstration that the between-state spatial variability in state-specific FC maps is significantly higher when modelled using the MAGE approach on UKB data compared with the HMM or SWC approaches. **[A-C]** The pair-wise Riemannian distance of the state-specific FC correlation matrices between 12 FC states is shown for MAGE, HMM and SWC, alongside the FC correlation maps for 5 of the identified states. The state-specific FC maps are shown as rank-one decomposition (first Eigenvector) of the estimated state-specific FC correlation matrices. The channels with positive values (red regions) show they are positively correlated with other regions, whereas channels with negative values (blue regions) show they are negatively correlated with other regions. The state-specific FC maps for all 12 states using SWC, HMM and MAGE are illustrated in Figure S14, Figure S15, and Figure S16 respectively. **[D]** The between-state Riemannian distance for the modelled 12 states, shown for MAGE, SWC and HMM. **[E]** Video for the time-point by time-point instantaneous FC estimation for a randomly chosen UKB subject is shown for the MAGE, SWC and HMM approaches.

To ensure that the MAGE estimated FC and mean activity states are reproducible, we ran MAGE six times on different non-overlapping subsets of UKB data (2k subjects for each run). In Figure 8[A,B], we compare the similarity between MAGE estimated state-specific spatial maps across the six runs. Specifically, in Figure 8[A], we show the average correlation plot for the state-specific FC estimates across repeated runs, and in Figure 8[B] for the state-specific mean activity estimates, both for a fixed number of states (P=Q=12). This shows that the state-specific FCs are very reproducible and that the state-specific mean activity estimates are moderately reproducible but not to the same extent. We then replicated these reproducibility findings for a wide range of numbers of states (1 to 24) for the FC and mean activity, as shown in Figure 8[C], also demonstrating that P does not have to be equal to Q. We show in Figure 8[C] that the FC are mostly highly and mean activity are mostly moderately reproducible independent of the number of modelled states. Specifically, mean activity and functional connectivity states are more reproducible if the number of modelled states is more than approximately six.

**Figure 8:**
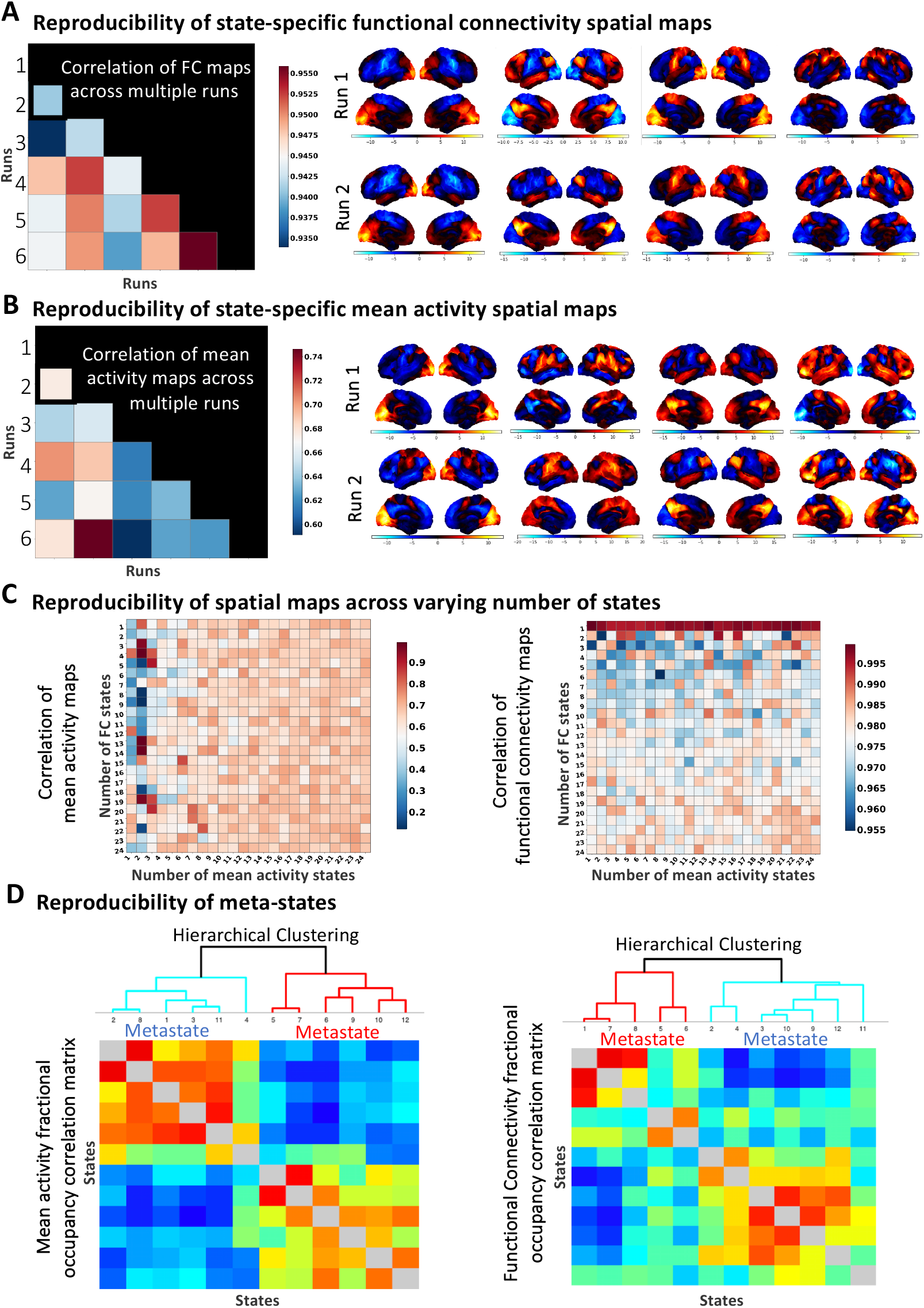
**[UKB data]** MAGE estimated FC and mean activity spatial maps are reproducible across various runs on different non-overlapping sets of subjects. **[A]** shows the average correlation between the state-specific FCs across six repeated runs. Moreover, for runs 1 and 2, FC correlation maps are shown for 4 of the identified states (out of the 12 inferred states), demonstrating that the FC states are consistent across repeated runs. On the other hand, **[B]** shows the average correlation plot for the state-specific mean activity estimates across six repeated runs, alongside the mean activity spatial maps (for runs 1 and 2 only). The analysis in **[A]** and **[B]** is performed for a fixed number of states (P = Q = 12), whereas, in **[C]**, we evaluated the reproducibility of state-specific FC and mean activity estimates by varying the total number of modelled states (P, Q). Specifically, **[C]** shows the correlation plots for the state-specific FCs and state-specific mean activity estimates, where we varied the total number of inferred states (P and Q) from 1 to 24. These results suggest that inferred states for the FC are highly and mean activity are moderately reproducible independent of the number of modelled states (Note the difference in the ranges of the colormaps). **[D]** We replicated the original findings of [Vidaurre et al. (2017)] and illustrate that hierarchical meta-state structure exists both in the functional connectivity and the mean activity states, but is more well-defined in the mean activity states (matrices computed as *vv*′, where *v* is the [number of states by number of subjects] fractional occupancy matrix).

It has been previously reported in [Vidaurre et al. (2017)] that the brain network activity is organized into a hierarchy of two distinct meta-states, with one meta-state representing hisgher-order cognition, and the other representing the sensorimotor systems. We have taken a similar approach as [Vidaurre et al. (2017)], and computed [P x d] or [Q x d] matrices containing the fractional occupancy (time spent in each state) for each subject, where d is the number of subjects. We then use these to compute the [P x P] or [Q x Q] correlation matrices over subjects, which are shown in Figure 8[D]. As in [Vidaurre et al. (2017)], Figure 8[D] shows that a hierarchical meta-state structure is present. Here, it is apparent in both in the functional connectivity and the mean activity driven state time-courses, although it is more apparent in the mean activity state time-courses.

### 3.6. Multi-dynamic TVFC predicts behavioural variability better than the single-dynamic approaches

We next used the UKB data to see if the multi-dynamic MAGE approach provides spatial network features (i.e., the state-specific FCs) that can be used to predict individual behavioural traits, and we compare its performance to the HMM and SWC approaches. Alongside the task fMRI results, this would provide further evidence that the state dynamics being inferred by MAGE are cognitively meaningful.

We applied the dual-estimation method to obtain subject-level estimates of TVFC. This means that for each subject, we can obtain a set of state-specific FC maps (Q * N * N), where Q is the number of correlation (FC) states and N is the number of channels/brain regions. Then, we can use these subject-level, state-specific FC estimates to predict selected behavioural variables. The classifier/predictor we employed is Elastic Net [Zou and Hastie (2005)], which is a regularized regression method that combines the penalties of lasso (*L*_1_) and ridge (*L*_2_) methods [Vidaurre et al. (2013)]. The data matrix fed into the classifier is d*[Q*f], where d is the number of subjects, Q is the number of states and f is the number of features. The state-specific FC matrices are symmetric and only values above the diagonal need to be retained and vectorized, and hence f = N(N-1)/2. Lastly, we concatenated the state-by-state subject-level, state-specific FC estimates, resulting in Q*f features for each subject. We performed this prediction analysis for UKB data with d = 13301, Q = 12, and N = 25.

We used four non-imaging variables to compare the performance of multi-dynamic (MAGE) approach with HMM and SWC. The details for the SWC and HMM approaches are explained in the Section S1.3 and Section S1.4 respectively. These variables are age, sex, fluid intelligence score, and neuroticism score. In Figure 9[A], we display the prediction results (nested 10-fold cross-validated) from the subject-level FC estimates. We can see that the FC obtained using the MAGE approach is a more accurate predictor of behaviour as compared to the other approaches

**Figure 9:**
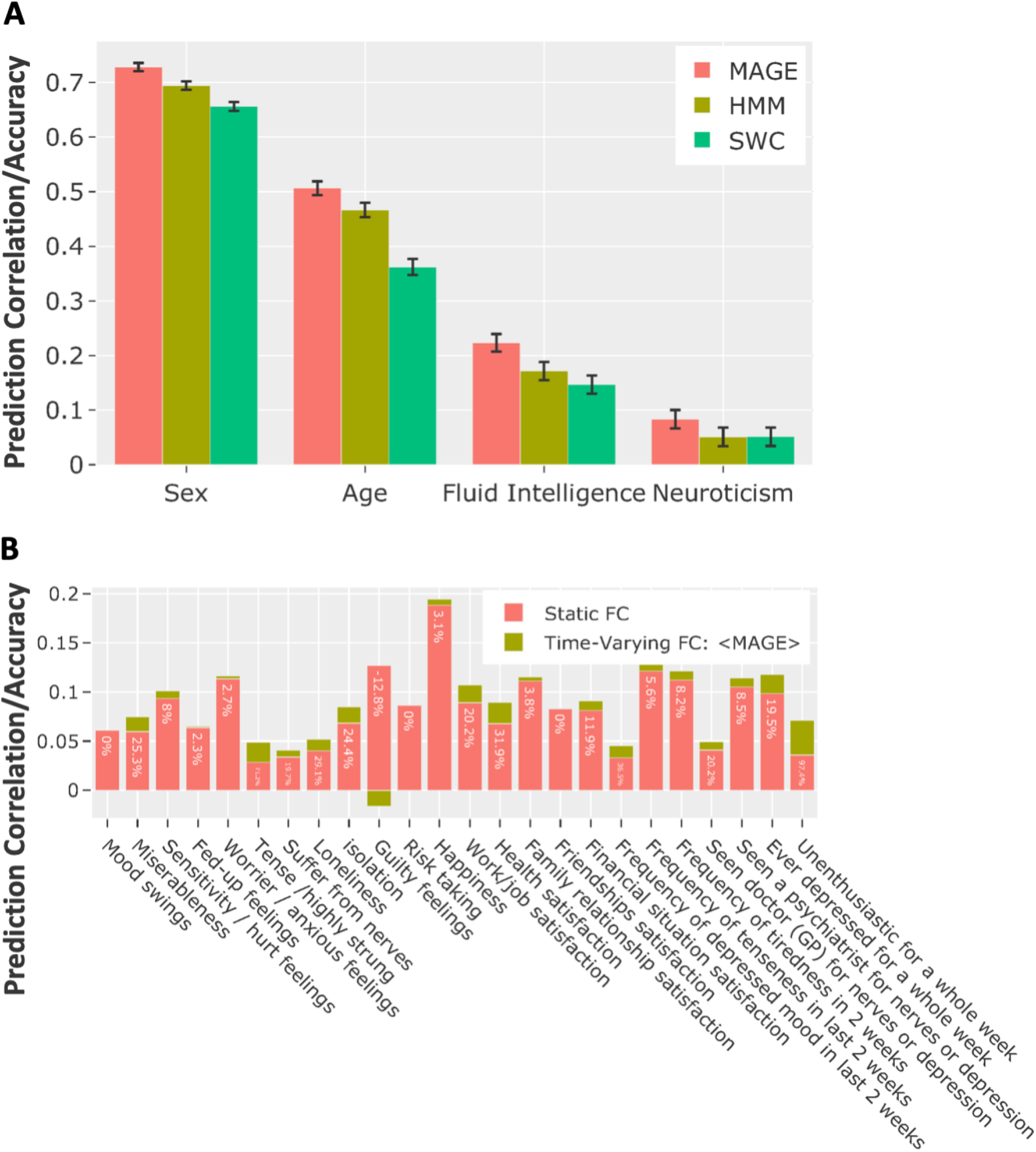
Multi-dynamic (MAGE) derived state-specific FCs are used to predict non-imaging variables in the UKB data. **[A]** The prediction performance of MAGE-derived state-specific FCs compared with the HMM and SWC approaches. The prediction accuracy/correlation is estimated using the combined features by concatenating the state-specific FC estimates from all states. Y-axis shows the prediction correlation (age, fluid intelligence, neuroticism) and prediction accuracy (sex), and the x-axis shows the predicted variable. To draw the confidence intervals on the prediction accuracies for sex variable (discrete output), we performed the Wilson test (suitable for the binomial distribution). Lastly, to generate confidence intervals on the prediction correlations (i.e., continuous outputs e.g., age, fluid intelligence and neuroticism), we computed the Fisher transformation [Pervaiz et al. (2020)]. **[B]** Comparison of the prediction performance of MAGE-derived state-specific FCs with the time-averaged FC. The bars in red colour show the prediction correlation/accuracy using time-averaged FC estimates, whereas bars in green show the increased/decreased prediction correlation/accuracy using state-specific FC estimates.

### 3.7. Static FC is worse than Multi-dynamic TVFC at predicting behaviour

Time-averaged FC (static) can be considered as an average of all state-specific FC states (and the time varying means) and is conventionally estimated simply by taking a full (Pearson) correlation between brain parcels over the whole scanning time. Unlike TVFC in the form of the state-specific FCs, time-averaged FC is a single FC estimate per subject and is already an established predictor of behavioural traits [Pervaiz et al. (2020)]. In Figure 9[B], we compare the combined power of the state-specific FCs with the time-averaged FC. This shows that for the mental health variables, predictive performance is improved by using the state-specific FCs in comparison to time-averaged FC. We employed Elastic Net for this comparison and the reported results are nested 10-fold cross-validated.

## 4. Discussion

We have proposed a novel algorithm, MAGE, that reliably infers the time-point by time-point estimates of FC. Notably, MAGE frees up the dynamics of the FC to fluctuate independently from the dynamics of the means and variances of the brain activity. We found that this multi-dynamic approach reveals much stronger changes in FC over time than existing single-dynamic approaches (e.g., HMM, SWC), and is a better predictor of individual behavioural variability. This provides a potential explanation and solution as to why time-varying FC has appeared to look so stable in resting fMRI data in previous work [Hindriks et al. (2016); Liegeois et al. (2017)].

### MAGE inferred multi-dynamic linear mixtures of FC states

We assessed the performance of MAGE on simulations where there can be more than one underlying state active at each time point (Figure 3). We demonstrated that categorical models, such as the HMM, under-performed as expected when applied to such simulations. In contrast, MAGE remained accurate across a range of these simulations. MAGE also shows improved performance in simulations designed to be suitable for categorical models (e.g., HMM), i.e., by having only one state active at each time point (mutual exclusivity) (Figure S3). This suggests that MAGE performance remained stable when the state time-courses have long-range temporal dependencies (i.e., violate the HMM Markovian assumption) (Figure S3[D]). Lastly, we assessed the performance on the multi-dynamic simulations and demonstrated that MAGE accurately identified unique dynamics of the correlations when the ground truth was that the correlations and mean activity dynamics were distinct from each other (Figure 4).

### MAGE learned network state dynamics that showed appropriate task dependencies

Following rigorous performance evaluation of MAGE on more than a thousand simulations, we applied it to a task fMRI dataset. The rationale was that task fMRI gives an opportunity to assess the specific state dynamics that MAGE infers, by comparing them to the known task timings. In Figure 5, we demonstrated that the FC-driven time courses performed well in terms of displaying task-dependencies as compared to the mean-driven state time courses when using MAGE. Moreover, we showed in Figure 5[B] that plausible time-varying representations of FC and mean activity spatial maps were inferred using the MAGE approach. This suggests that the inferred TVFC is powerful enough to capture the dynamics in cognitive tasks, and is not static during these task experiments.

### Why do we model the mean of the brain activity separately from the functional connectivity?

Our assumption in Equation 1 is that the mean and the FC (i.e., the correlation matrix) continuously vary over time. From that perspective, there is no imperative to assume that the dynamics of those two entities are the same as each other. As such, MAGE is designed to explore what happens when we do not tie the dynamics together. By allowing the state dynamics of the mean and FC to be independent of each other, MAGE also allows them to potentially operate on different time scales. However, in real fMRI data, we found that the dynamics of the mean states and the FC states ended up on fairly similar time-scales (between 2 to 5 seconds for FC states and 2 to 3 seconds for mean activity states, as shown in Figure 6). It is also interesting to consider the extent to which the state dynamics of the mean and FC get inferred as being coupled together, despite being free to be inferred as completely independent. Figure 6[C] demonstrated that there was indeed some significant non-zero correlation between the two types of dynamics. However, this correspondence between the dynamics of the mean and the dynamics of the FC was not absolute. In other words, the fluctuations in the FC clearly have some unique temporal structure that is not shared with the fluctuations in the mean of the activity. Indeed, the benefit of capturing these unique fluctuations was demonstrated by the improved performance of MAGE (versus SAGE) in inferring stronger fluctuations in FC over time and in predicting behavioural traits.

### Why does MAGE reveal stronger functional connectivity dynamics in resting-state fMRI?

Our results demonstrated that MAGE revealed stronger changes in FC over time than single-dynamic approaches, where changes in FC over time correspond to between-state differences in the state-specific FCs (e.g., HMM, SWC, SAGE, see Figure 6 and Figure 7). The lack of change in FC over time in single-dynamic approaches has been observed in other, recent studies [Laumann et al. (2017); Hindriks et al. (2016); Leonardi and Van De Ville (2015)]. Indeed, in some cases, it has been observed that the reported variability in the FC from one state to another state (or against null models) is statistically insignificant [Liegeois et al. (2017)]. So why does MAGE show comparatively stronger changes in FC over time? We can gain insight into the multi-dynamic model being used in MAGE, by considering a dynamic version of structural equation modelling (SEM), as explained in Appendix A. This shows that if the underlying truth is that there are distinct dynamics for the external driving inputs and connectivity, then distinct dynamics are needed for the mean activity and FC modelled by MAGE. If we were to ignore this, and use a single dynamic approach (e.g., HMM, SAGE, SWC) then we will infer common state dynamics that are a compromise. In turn, use of these incorrect state dynamics will tend to cause the inferred FC patterns to be temporally smeared versions of the true FC patterns, causing them to look more homogeneous over time. We demonstrated this phenomenon with a simple example (Figure 1[B]), in which we simulated a ground truth where the FC dynamics and the mean activity dynamics were not tied together. Using SAGE on this data substantially reduced the inferred change in instantaneous FC over time, whereas MAGE better captured the true variability in instantaneous FC across time. This work provides a potential explanation as to why FC in resting fMRI has previously appeared to look so homogeneous over time [Laumann et al. (2017); Hindriks et al. (2016); Liegeois et al. (2017)].

### Reproducibility of MAGE states

The presented results in Figure 8 demonstrated that the MAGE estimated FC states are highly and mean activity states are moderately reproducible across re-runs on non-overlapping subsets of the data (shown quantitatively and qualitatively in Figure 8[A,B]). Also, as is typical with other unsupervised learning methods (e.g., ICA, HMM), the number of states to be modelled is an empirical parameter in our method and it needs to be chosen before running the model. In most of our presented work, we modelled anywhere from 4 to 16 states, though we had experimented with up-to 64 states in the resting-state and task data (and the results remained sensible). In Figure 8[C], we found out the estimated states are more consistent across re-runs when the number of modelled states is more than six. Considering that, we suggest that the number of modelled states to be more than approximately six for such dynamic modelling (though we do not have a strong recommendation for an exact number of states).

### Multi-dynamic TVFC predicted behavioural variability better than existing methods

We evaluated whether the inter-individual differences inferred by MAGE are meaningful. We demonstrated in Figure 9[A] that the MAGE-estimated, state-specific FCs did a better job at predicting these non-imaging variables as compared to SWC and HMM approaches. For example, age was predicted with an average correlation of 0.50 using MAGE as compared to 0.46 using HMM, and 0.36 using SWC. This provides evidence that the TVFC estimated using the MAGE is a more accurate and meaningful representation of underlying functional networks and is indeed a powerful biomarker for cognitive traits.

### Static FC is worse than Multi-dynamic TVFC at predicting behaviour

In Figure 9[B], we compared the predictive power of MAGE-derived, state-specific FCs versus time-averaged FC, and demonstrated the prediction performance on mental-health related variables in UKB data. We have demonstrated that the prediction accuracy of these behavioural variables increased when TVFC was used, as compared to time-averaged FC e.g., for work/job satisfaction, prediction accuracy was improved by 20.2 % and for depressed for a week, accuracy was improved by 19.5 %, etc. This highlights that there are some aspects of behaviour that can be best explained using TVFC estimates.

### What do we mean by TVFC?

We can gain insight into the multi-dynamic model being proposed, by considering a dynamic version of structural equation modelling (SEM) as explained in Appendix A. The SEM perspective (Appendix A) suggests that the instantaneous FC is the closest representation that we have to the time-varying connectivity in a dynamic SEM (Equation 19). In MAGE, we have assumed that the instantaneous FC, *C*_*t*_, is the TVFC measure of most interest. MAGE provides a regularised estimate of the instantaneous FC, and so in this paper we have represented the instantaneous FC parsimoniously using the state-specific FCs. Hence, in this work we have focussed on the state-specific FCs as our main measure of TVFC, e.g., for predicting behavioural traits. However, a more complete representation of TVFC includes the FC state time courses alongside the state-specific FCs (after all, we combine the state-specific FCs with the FC state time courses via Equation 4 to get the instantaneous FC). As such, prediction may be improved further in the future by including features describing the FC state dynamics. For example, we can already show that inter-individual differences for the time spent in each FC state are linked to behavioural traits, as illustrated in Figure S8. MAGE also provides other potential features for use in prediction, for example, the mean-activity FC (i.e., correlations of the mean activity, *m*_*t*_, over all time) or the state-specific mean activity and the mean-derived dynamics.

### Variance of the brain activity

We found through resting-state and task data-sets results (Figure 6 and Figure 7) that there was not much between-state variability in the variance of the activity, and the variance remained static over the scanning session. Nevertheless, we explored another configuration where dynamics of the variance of the activity was modelled with independent state dynamics from the mean activity and the FC, but that also did not result in any meaningful variability in the variance of the activity (though it is computationally more expensive to learn the variance state time courses separately). We also attempted to model state dynamics of the variance of the activity tied to the state dynamics of the FC fluctuations, but independently from the mean activity; however, this resulted in comparatively less between-state variability in the FC. In summary, we found out there was little between-state variability in the state-specific variances irrespective of the modelling choice.

In summary, we proposed a multi-dynamic approach (MAGE) that models temporal fluctuations in functional connectivity independently from fluctuations in the mean of the activity. Multi-dynamic modelling provided an explanation and a solution as to why resting fMRI FC has previously looked so stable and homogeneous. We found out that that separating fluctuations in the mean activity levels from those in the functional connectivity reveals much stronger changes in functional connectivity over time. Lastly, MAGE estimated time-varying FC is a better predictor of behavioural variability in the resting-state fMRI data than established methods.

## 5. Credit authorship contribution statement

**Usama Pervaiz:** Data curation, Conceptualization, Methodology, Writing - original draft, Validation, Visualization. **Diego Vidaurre:** Methodology, Writing - review and editing, Supervision. **Chetan Gohil:** Methodology, Writing - review and editing. **Stephen M. Smith:** Data curation, Conceptualization, Writing - review and editing, Validation, Supervision. **Mark W. Woolrich:** Conceptualization, Methodology, Writing - original draft, Writing - review and editing, Validation, Visualization, Supervision.

## 6. Acknowledgements

Computation used the Oxford Biomedical Research Computing (BMRC) facility, a joint development between the Wellcome Centre for Human Genetics and the Big Data Institute supported by Health Data Research UK and the NIHR Oxford Biomedical Research Centre.

We are extremely grateful to all UK Biobank (Data Access Application 8107) and Human Connectome Project participants who generously donated their time to make this resource possible. UK Biobank (including the imaging enhancement) has been generously supported by the UK Medical Research Council and the Wellcome Trust. Human Connectome Project, WU-Minn Consortium (Principal Investigators: David Van Essen and Kamil Ugurbil; 1U54MH091657) funded by the 16 NIH Institutes and Centers that support the NIH Blueprint for Neuroscience Research; and by the McDonnell Center for Systems Neuroscience at Washington University.

We additionally thanks Eugene Duff for providing the data access to the cognitive task fMRI data and Samuel Harrison for pre-processing the task data. Furthermore, we are thankful to Ryan Timms and Evan Roberts for the helpful discussions.

MWW’s research is supported by the NIHR Oxford Health Biomedical Research Centre, the Wellcome Trust (106183/Z/14/Z, 215573/Z/19/Z), the New Therapeutics in Alzheimer’s Diseases (NTAD) study supported by UK MRC and the Dementia Platform UK (RG94383/RG89702) and the EU-project euSNN (MSCA-ITN H2020-860563). The Wellcome Centre for Integrative Neuroimaging is supported by core funding from the Wellcome Trust (203139/Z/16/Z).

UP’s research is funded by an MRC Mental Health Data Pathfinder award (PI Clare Mackay) MC/PC/17215. SMS’s research is supported by the XMI2: Wellcome Trust Collaborative Award 215573/Z/19/Z. DV’s research is supported by Novo Nordisk Foundation Emerging Investigator Fellowship (NNF19OC-0054895) and ERC Starting Grant (ERC-StG-2019-850404).

## 7. Declaration of interests

The authors declare that they have no known competing financial interests or personal relationships that could have appeared to influence the work reported in this paper.

## Appendices

### A. Dynamic Structural Equation Modelling (SEM) perspective

We can gain insight into the multi-dynamic model being proposed, by considering a dynamic version of structural equation modelling (SEM) [Penny et al. (2004)]:

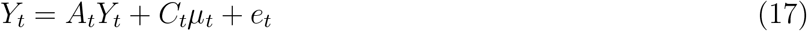

where *A*_*t*_ is an N*N dimensional matrix containing the connectivity information, where *A*_*ij*_ denotes a connection from region j to region i and *A*_*ii*_ = 0 for all regions (in SEM, *A*_*t*_ is referred to as the path coefficient matrix, M). *µ*_*t*_ is an Qx1 vector of Q structured, external inputs that are independent such that 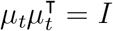, and *C*_*t*_ is an NxQ dimensional mixing matrix describing how much of each external input is contributing to each region. Finally, *e*_*t*_ is a random variable modelling the noise in *Y*_*t*_ using a Gaussian distribution with zero mean and an NxN diagonal covariance matrix, i.e., *e*_*t*_ ∼ 𝒩 (0, Σ). Note that the idea is that *µ*_*t*_ would be identifiable through its non-Gaussian, spatio-temporal structure, which is what makes it distinct to *e*_*t*_. Note that typically in SEM, only the noise, *e*_*t*_, is modelled and an external input, *µ*_*t*_, is not (i.e. *µ*_*t*_ = 0). However, other variants of SEM have previously considered modelling structured, external inputs [Shimizu et al. (2006)].

It is straightforward^5^ to then show that *Y*_*t*_ is Gaussian distributed, with a time-varying mean and covariance with the same form as in Equations (3, 4), where

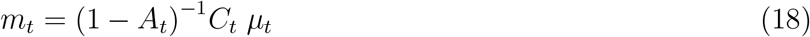

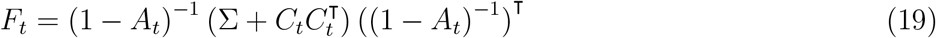

In other words, if we assume that in the dynamic SEM the external inputs, *µ*_*t*_, and the dynamic connectivity, *A*_*t*_, fluctuate distinctly from each other; then correspondingly *m*_*t*_ and *F*_*t*_ can also fluctuate distinctly from each other (see Equation 18, 19) also need to fluctuate distinctly from each other. This is what we are assuming in the *multi-dynamic* approach.

## Footnotes

2 The reason MAGE models the variances to be independent of the correlation (i.e., *γ*_*tr*_ free to be not equal to *β*_*tq*_) is to allow maximum temporal independence to the correlation. We will expand onto this in Section 4 but allowing *γ*_*tr*_ = *β*_*tq*_ resulted in weaker state-specific FCs dynamics, and hence was not a preferable modelling choice.

3 For this particular simulation, the number of channels in the data is 20, the number of modelled states (P=Q=R) is 12, and multi-dynamics state time courses for the means and the correlations are simulated using two different hidden semi-Markov model chains.

4 For other such examples, see [Vidaurre et al. (2021)] for modelling only the covariance and [Charquero-Ballester et al. (2020)] for modelling only the mean activity.

5 Such investigations to establish the relationship between SEM and FC has been previously conducted in detailed by [Marrelec and Benali (2009)]

## Notes

### Competing Interest Statement

The authors have declared no competing interest.

