## Supplementary File for "Multi-dynamic Modelling Reveals Strongly Time-varying Resting fMRI Correlations"

### S1. Method's Supplementary

#### *S1.1. Single-dynamic (SAGE) Model*

SAGE model assumes that the mean, variance, and correlation fluctuate on a single (common) time-course. This implies that the time courses $\alpha_{tp}$, $\beta_{tq}$ and $\gamma_{tr}$ are the same in the SAGE approach. The time-varying mean, correlation and variance are modelled as:

$$m_t = \sum_p \alpha_{tp} \mathcal{S}_p \tag{S1}$$

$$F_t = \sum_p \alpha_{tp} \mathcal{D}_p \tag{S2}$$

$$G_t = \sum_p \alpha_{tp} \text{diag}(\mathcal{E}_p) \tag{S3}$$

where, the $\alpha_t$ is modelled by underlying logits $\theta_t$. The loss function for SAGE is:

$$\sum_{t=1}^{T} log\, p(Y_t|\theta_t = \mu_t^\theta) + \sum_{t=1}^{T} \text{CrossEntropy}\,(\mu_t^\theta || \hat{\mu}_t^\theta) \tag{S4}$$

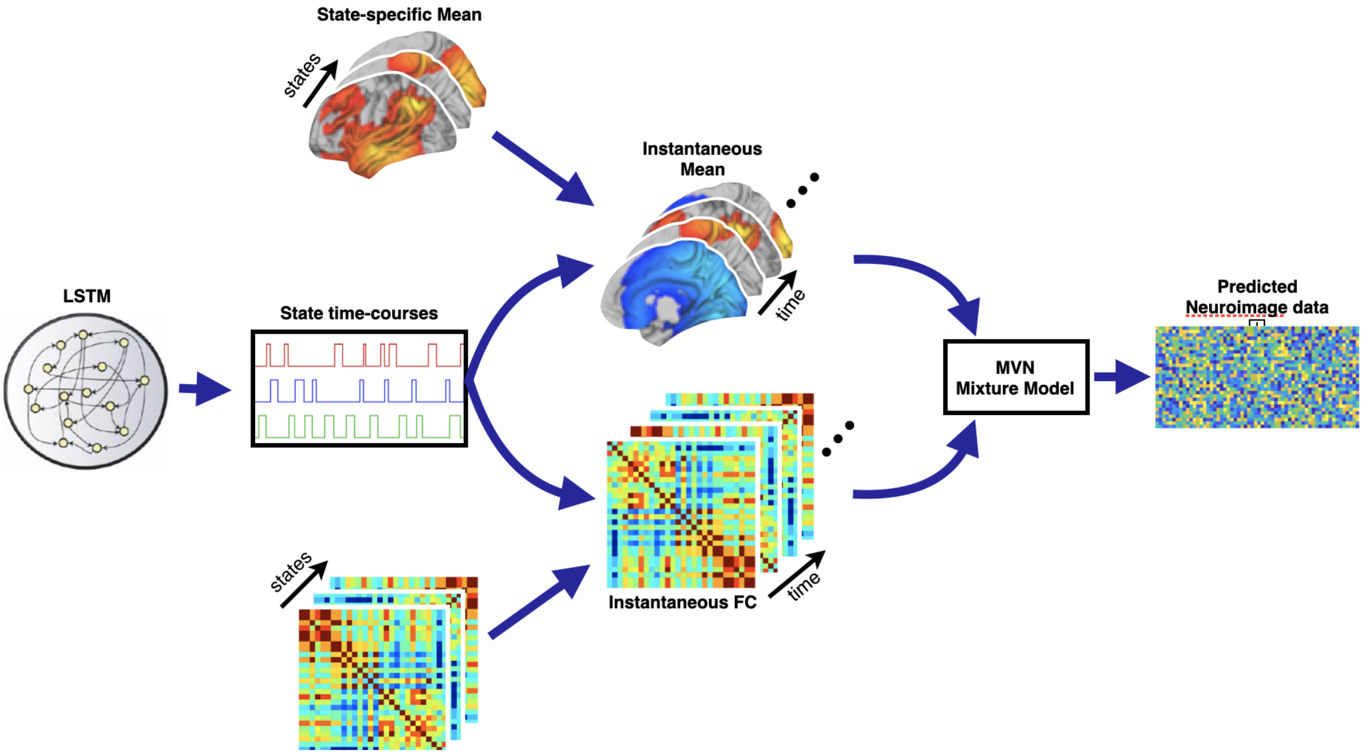


Figure S1: Generative model for SAGE. The instantaneous mean and correlation are modelled using an underlying set of states, for which the state time courses are generated using a long short-term memory (LSTM) model (common for both the mean and correlation), making the approach single-dynamic.

*S1.2.* ***MAGE In-depth architecture***

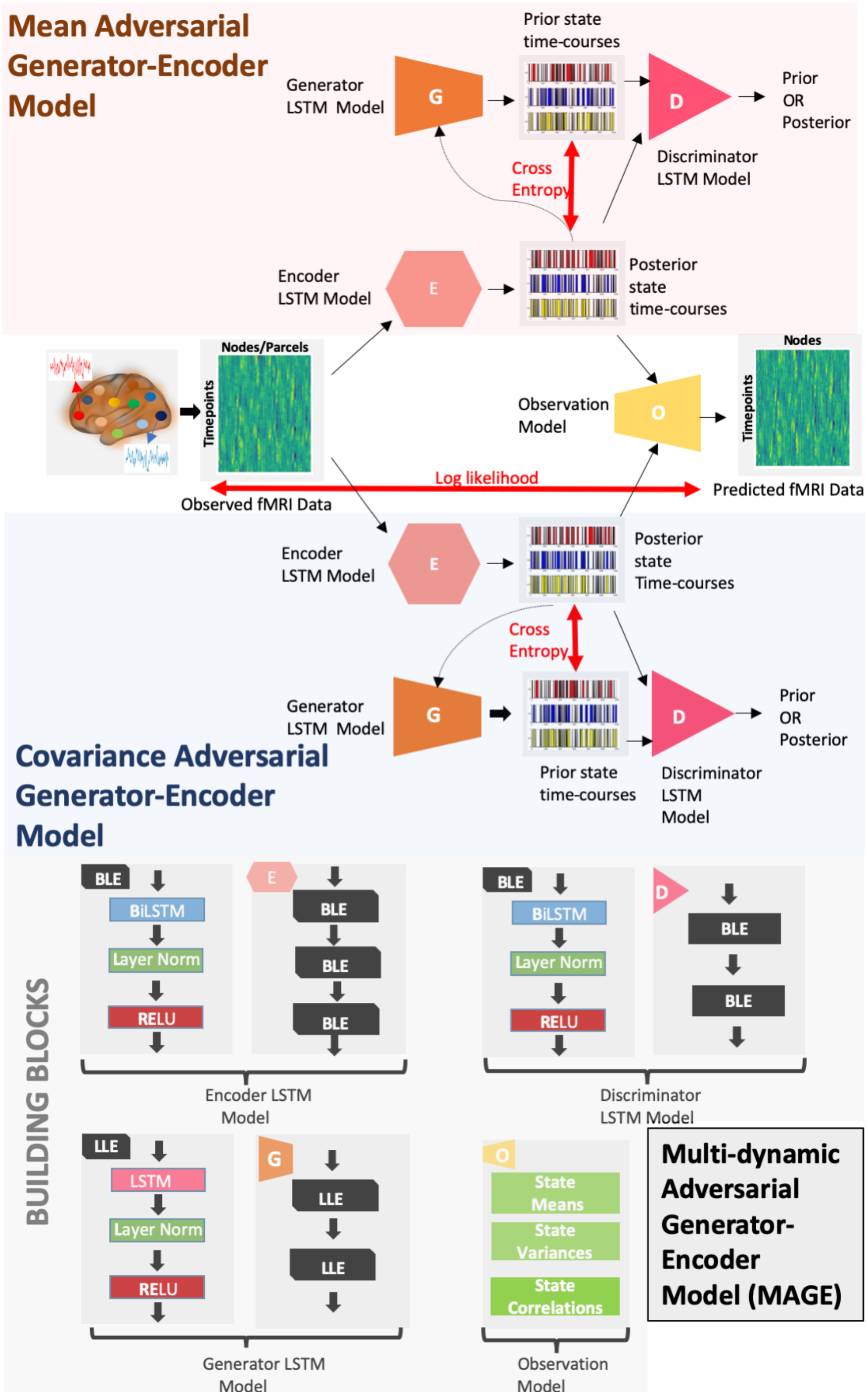


Figure S2: The in-depth architecture of the MAGE is illustrated here. For the multi-time-scale modelling, two sub-blocks are proposed; one for the generation-inference of covariance associated parameters and the other for the mean. Each block is composed of a generator, encoder, and discriminator model. The generator model predicts the prior state time-courses and the encoder model predicts the posterior state time-courses, whereas, the discriminator model enforces the prior state time-courses on the posterior state time-courses. The encoder model is composed of three BLE blocks, whereas each BLE block is composed of a bi-directional LSTM, layer normalization, and a rectified linear activation function (ReLU) layer. The discriminator is composed of two BLE blocks whereas, the generator model is composed of two LLE blocks. The main difference between LLE and BLE block is the usage of a uni-directional LSTM instead of a bi-directional LSTM.

#### *S1.3.* ***Sliding Window Correlation (SWC) approach***

For the SWC analysis, given a window of length, w, we removed all the frequency components below 1/w. Specifically, we have chosen a window-length of 100 seconds and estimated pair-wise correlations within each window, resulting in one FC correlation matrix per window. We used overlapping windows with an overlap of 25 time-points between two adjacent windows. After performing the SWC analysis, we applied the k-means clustering to the SWC estimated FC matrices. This clustering was applied at the group-level so the identified states are comparable across subjects. We associated each FC matrix with a clustering label/state, whereas, the total number of k-means states, are set to 12 (same as the MAGE and HMM approach). Then, we averaged the FC matrices for each subject as per the clustering labels. This sliding window *plus* clustering analysis provided us with TVFC estimates for each subject where the maximum number of states for each subject is 12 (and a state is not included if it is not active for that subject). Lastly, we concatenated the state-by-state FC matrices for each subject for any post-hoc prediction analysis conducted in the paper.

#### *S1.4.* ***Hidden Markov Modelling (HMM) approach***

For the HMM analysis conducted in this paper, states are modelled using multivariate Gaussian distributions, each state has a distinct mean and a distinct covariance matrix. In the HMM, the parameters describing the dynamics are categorical state time courses. Specifically, we model the covariance $C_t$ at each time point as a selection of a single covariance matrix from a finite set of covariances, $\mathcal{D}_p$ where p = [1...P]. Similarly, we model the mean $m_t$ at each time point as a selection of a single mean vector from a finite set of mean vectors, $\mathcal{S}_p$.

The mean and covariance at any time-point, t, can be modelled as:

$$C_t = \mathcal{D}_p, m_t = \mathcal{S}_p$$

where, $\mathcal{D}_p$ and $\mathcal{S}_p$ depends on the state active at each time-point, t:

$$\alpha_t = p \tag{S5}$$

where, $\alpha_t$ is a state time course indicating which state (out of P possible states) is active at each time-point, t.

If $Y_t$ represents the data, and $\alpha_t$ represents the hidden state at time point t, then the HMM assumes that:

$$Y_t | \alpha_t = p \sim \mathcal{N}(m_t, C_t) \tag{S6}$$

#### *S1.5.* ***Derivation of the regularization loss function for the SAGE approach***

The loss function for the SAGE approach is stated in Equation S4. The second term of the loss function is the regularization term that minimises the cross-entropy between the prior and the posterior of the latent, time-varying state time-courses.

$$\sum_{t=1}^{T} \text{CrossEntropy}\left(\mu_t^\theta || \hat{\mu}_t^\theta\right) \tag{S7}$$

To achieve this, proposed loss function is composed of two terms, the encoder adversarial loss and the generator adversarial loss.

***Encoder regularization loss***. We trained the discriminator (D) model to distinguish posterior from the prior. Simultaneously, we modified the encoder (E) model to make the discriminator believe that encoder model outputs are samples from the generator (G) model.

The objective of the discriminator model is to correctly classify the $\mu_t^\theta$ from $\hat{\mu}_t^\theta$. For this, Equation S8 should be maximized (max) and the loss function for the discriminator model can be written as:

$$\sum_{t=1}^{T} \underset{D}{max}\ [logD(G(\hat{\mu}_t^\theta))] +\ [log(1 - D(E(\mu_t^\theta))] \tag{S8}$$

On the other hand, we make the encoder model compete against the discriminator model, and it would try to minimize (min) Equation S8 as shown below:

$$\sum_{t=1}^{T} \underset{E}{min}\ [logD(G(\hat{\mu}_t^\theta)] +\ [log(1 - D(E(\mu_t^\theta))] \tag{S9}$$

If we combine the Equation S8 and Equation S9, It can be written as:

$$\sum_{t=1}^{T} \underset{D}{max}\ \underset{E}{min}\ [logD(G(\hat{\mu}_t^\theta))] +\ [log(1 - D(E(\mu_t^\theta)))] \tag{S10}$$

In summary, the encoder regularization loss has two objectives; the objective of the discriminator model is to learn to distinguish posterior from the prior, and the objective of the encoder model is to fool the discriminator model into believing that $\mu_t^\theta$ is not inferred from the encoder model but the generator model.

***Generator regularization loss***. For the generator adversarial loss, we again trained the discriminator model to distinguish posterior from the prior. Unlike the encoder adversarial loss, this time we trained the generator model to make the discriminator believe that generator model outputs are samples from the encoder model.

The objective of the discriminator is:

$$\sum_{t=1}^{T} \underset{D}{max}\ [logD(E(\mu_t^\theta))] +\ [log(1 - D(G(\hat{\mu}_t^\theta))] \tag{S11}$$

The generator model here would be competing against the discriminator model and would try to minimize Equation S11 as:

$$\sum_{t=1}^{T} \underset{G}{min}\ [logD(E(\mu_t^\theta))] +\ [log(1 - D(G(\hat{\mu}_t^\theta))] \tag{S12}$$

Equation S11 and Equation S12 can be written in condensed form as:

$$\sum_{t=1}^{T} \underset{D}{max}\ \underset{G}{min}\ [logD(E(\mu_t^\theta)] +\ [log(1 - D(G(\hat{\mu}_t^\theta))] \tag{S13}$$

In summary, the generator regularization loss has two objectives, the objective of the discriminator is to learn to distinguish posterior from the prior, and the objective of the generator is to fool the discriminator model into believing that $\hat{\mu}_t^\theta$ is not inferred from the generator model but the encoder model.

***Total regularization loss.*** The total adversarial loss can be written by combining Equation S10 and Equation S13:

$$\sum_{t=1}^{T} \underset{D}{max}\ \underset{G}{min}\ \underset{E}{min}\ [logD(G(\hat{\mu}_t^\theta)] +\ [logD(E(\mu_t^\theta))] +\ [log(1 - D(E(\mu_t^\theta))] +\ [log(1 - D(G(\hat{\mu}_t^\theta))] \qquad \text{(S14)}$$

This derivation can be easily extended to the multi-dynamic (MAGE) approach, where we are modelling $\theta_t^m$ and $\theta_t^c$ instead of a single $\theta_t$.

### S2. Result's Supplementary

#### *S2.1.* ***MAGE Performance on HMM-like Categorical Simulations***

We generated a multivariate normal distribution based on the assumptions that the state time courses are categorical and then simulated a multi-channel data-set from the distribution. For the simulation of the state time courses, we used Markov and semi-Markov chains, where transitions are dictated by a transition probability matrix (the probability of transition from one state to another). The main difference between the semi-Markov chain and the Markov chain is that the state lifetimes (i.e. how long one state will stay active before transitioning to a new state) in the semi-Markov chain are not governed by a Markovian dependency. Instead, they are, for example, sampled from a distribution of state lifetimes. The rationale behind the Markov chain based simulations is to match the HMM assumptions, and to demonstrate that MAGE performs well even in those scenarios. Moreover, the reason for including the semi-Markov chain based simulations is to determine MAGE performance when the state time-courses have long-range temporal dependencies (i.e., violate the HMM Markovian assumption).

We have illustrated the performance of MAGE across several categorical scenarios in Figure S3 (and compared the performance with the HMM and SWC). These simulated scenarios have certain hyper-parameters in common, e.g.,: 1) The number of channels, N, in the data. 2) The number of modelled states, P and Q and R. 3) The percentage of observational error, E. 4) Duration of the state lifetimes, L, for semi-Markov chains. 5) The number of batches, B, used for training the data. The default values (unless specified otherwise) in these simulations are: N = 20 channels, P = Q = R = 6 states, E = 0.04% and B = 200 batches. For each of the simulation configurations, we generated fifty different simulations, in which we varied the observation model (randomly generated mean and covariance) and the state time courses (with randomly generated transition probability).

The performance is assessed by measuring the accuracy between the simulated versus predicted states using Markov chains in Figure S3[A-C]. In Figure S3[A], N is varied from 20 to 100, whereas, in Figure S3[B], P, Q and R are varied from 2 to 16 and in Figure S3[C], E is varied from 0.02% to 0.2%. Each violin plot shows the prediction variability over 50 different simulations, and MAGE performance is compared with HMM and SWC in all the above-mentioned scenarios. In Figure S3[D,E], we conducted the stability analysis of the proposed MAGE model based on two factors: 1) How the performance is changed as we increase the lifetime values of state time courses, and 2) How much data is required to train the model to obtain a satisfactory performance? For Figure S3[D], we simulated semi-Markov chains that are characterized by L (varied from 1 to 400 time points).

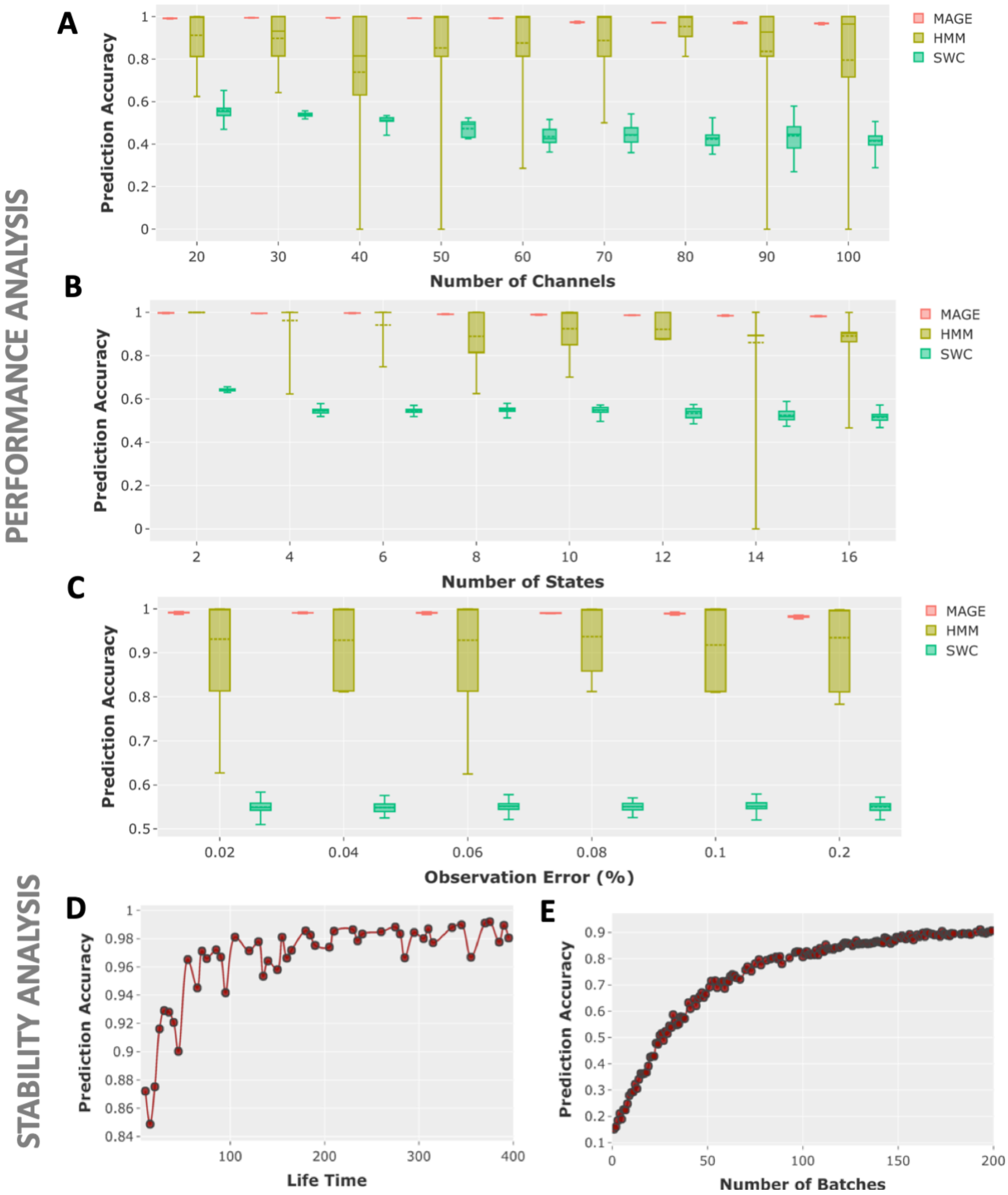


Figure S3: The performance and stability analysis of MAGE on the categorical simulations. MAGE, Sliding Window Correlation (SWC) and the Hidden Markov Model (HMM) were compared using same simulation parameters. In [**A-C**], the impact of varying the simulation parameters on the prediction accuracy of the MAGE, HMM and SWC was assessed using Markov chains. Each violin plot shows the prediction variability over 50 different simulations. Specifically, we have illustrated the performance as we varied number of channels in [**A**], the number of states in [**B**], and observational error percentage in [**C**]. In [**D-E**], we conducted a stability analysis to see the impact on performance of MAGE as we increased the ground truth lifetime of the simulated states using semi-Markov chains [**D**] or as we increased the number of batches [**E**].

#### *S2.2.* ***MAGE performance on static-FC simulations***

We demonstrated the performance of MAGE for scenarios where the ground-truth is such that correlation (i.e., the FC) is static (i.e., no fluctuations in the edges of the FC matrix over time). We show in Figure S4 that MAGE does not infer time-varying correlations when we infer more than one state for such simulations. It highlights that MAGE only infers time-varying FC when the underlying FC is time-varying.

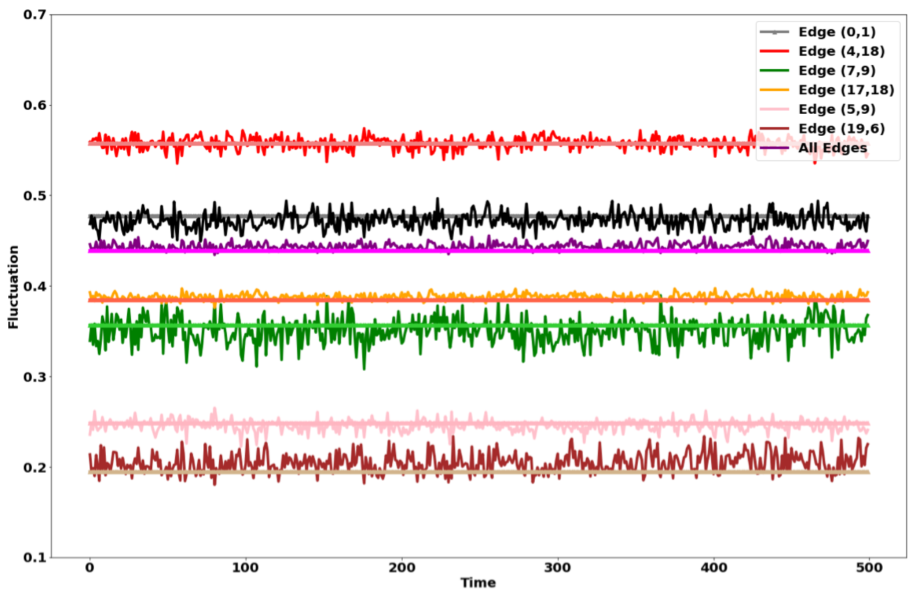


Figure S4: MAGE does not infer time-varying FC when the underlying ground-truth is not time-varying. The ground-truth is that there is only one state active at all the time-points (implies that the FC is not fluctuating). In such a simulation, even when we infer more than one state, MAGE only infers the static FC (closer to the ground-truth). We have illustrated such results for six edges and average across all the edges of the FC matrix. The light-faced straight lines highlight the ground-truth that shows the instantaneous FC is not fluctuating across time for these six edges. The bold-faces lines around the ground-truth highlight the infer instantaneous FC estimates also look homogeneous across time.

#### *S2.3.* ***MAGE performance on fluctuating mean activity and null (static) HCP data***

We generated null data using autoregressive randomization (ARR). Specifically, a 10-th order Gaussian autoregressive (AR) model was applied to generate null data. For each participant and each pair of brain regions, we compared the observed between-timepoint variances of the instantaneous FCs inferred by MAGE, which can be calculated from the FC state dynamics and state-specific FCs, from the real HCP data against the null data. This analysis resulted in one p-value for each edge of the FC matrix. On average, across HCP participants (compared against null data), 95% of edges were significant ($p \leq 0.001$).

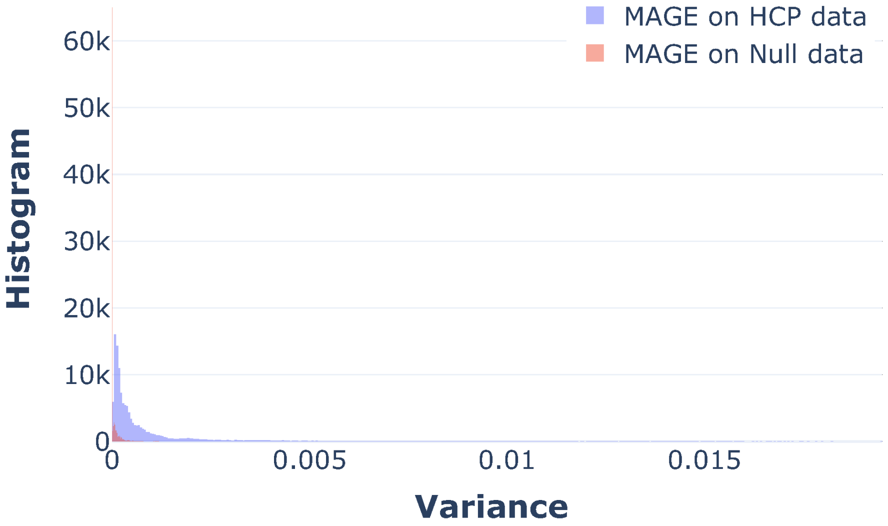


Figure S5: MAGE results on null (static) HCP data. The histogram of null variance estimates is compared with the variance estimates from the real HCP data for each participant and each pair of brain regions. The histogram of the variance estimates highlights that the between-brain region variability over time is significantly higher in the real HCP data as compared to null HCP data.

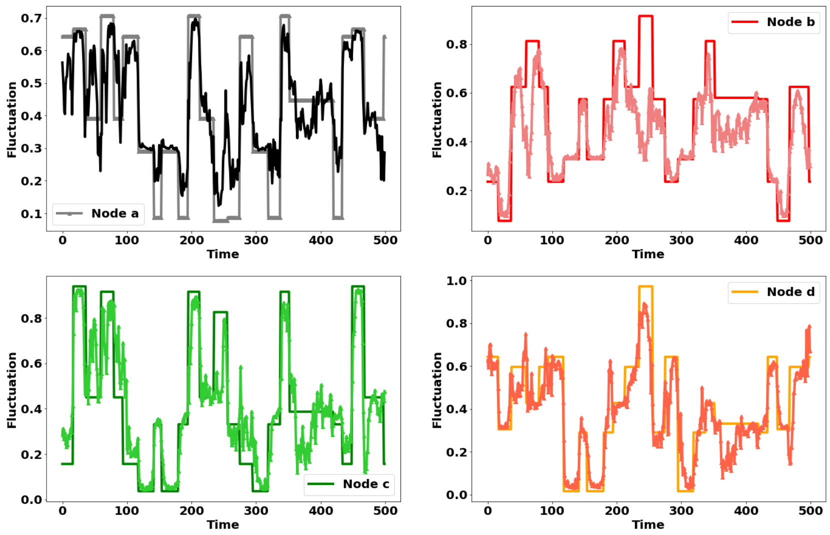


Figure S6: MAGE can adjust to fluctuating mean activity. It is the continuum of the multi-dynamic results presented in Figure 1 of the article. The multi-dynamic approach can adjust to fluctuating FC (as shown in Figure 1) and also accurately adjust to fluctuating mean activity of nodes (as shown here). We illustrated these results for four of the nodes of the FC matrix where we have shown the ground-truth and the inferred estimates.

#### *S2.4. **Task Data Results on 8 and 16 states***

We have illustrated the results of task decoding when we modelled more than 4 states by linking the fractional occupancy (the time spent in each state) with the experimental task. In Figure S7[A] and Figure S7[B], we have modelled 8 and 16 states respectively. Particularly, we plotted the relative fractional occupancy (mean change in occupancy relative to baseline) distribution for each of the modelled states. It is evident that even with the increased number of states, we can still find task-specific states (as highlighted by the paired t-test values on the respective states).

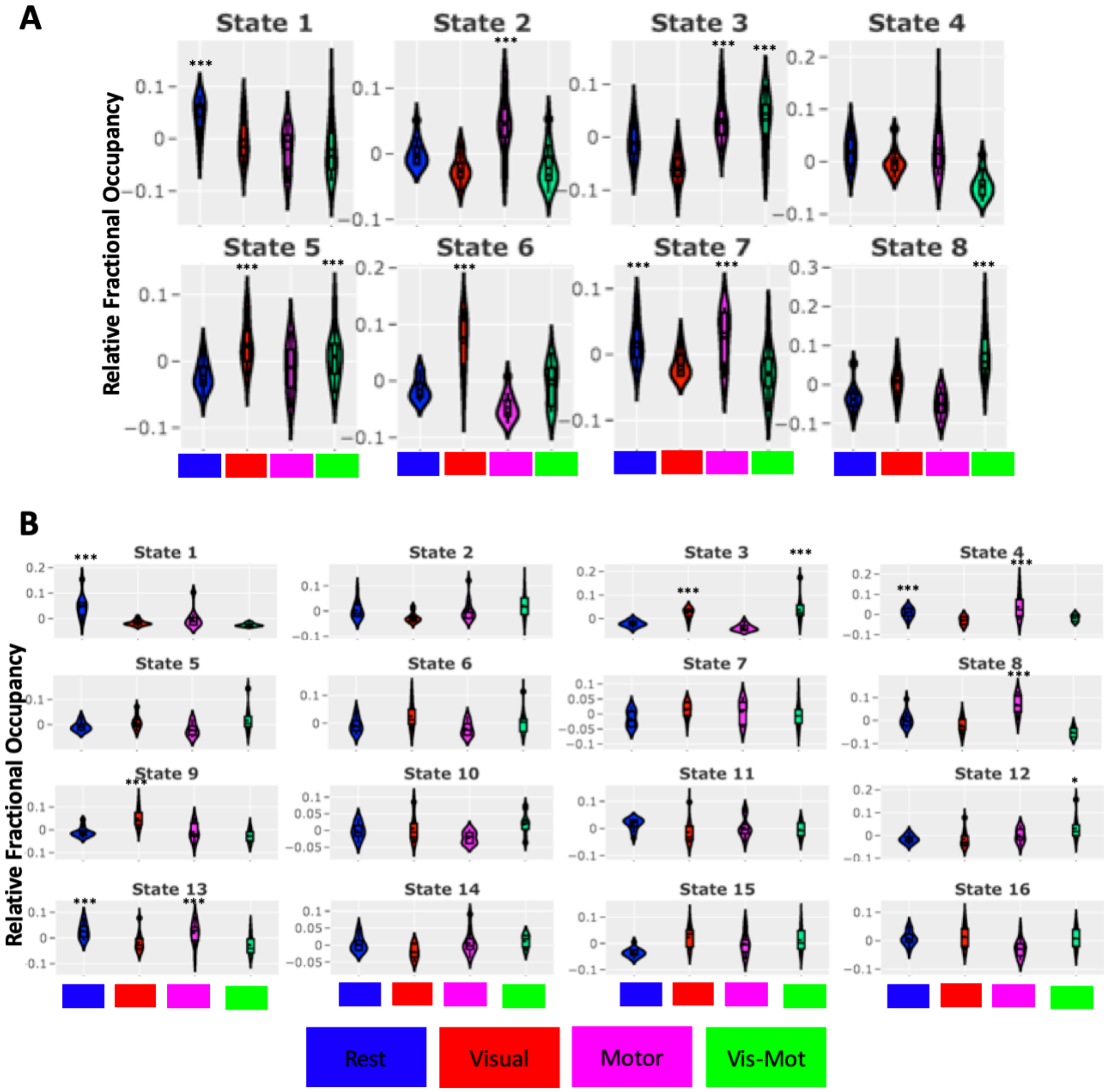


Figure S7: [**A-B**] We illustrated the results of task decoding for the task data set when more than 4 states are modelled. On the y-axis, we have shown the relative fractional occupancy response for each of the modelled states, and the on x-axis, we have highlighted the experimental task at hand. The paired t-test highlights the correspondence of states with the experimental task: [**A**] State 1 is mostly active during the rest, state 2 is active during the motor task, state 3 is active for both motor and visual-motor task, and state 8 is dominantly active for the visual motor task, etc. [**B**] State 1 and State 13 is active during the rest, state 3 and state 9 is active during the visual task, etc.

*S2.5.* ***Rest Data Results: Correlation with behaviour using Fractional Occupancy***

The goal here is to specifically look into the FC-driven state time courses and achieve a more meaningful understanding of FC temporal dynamics. We assessed how strongly fractional occupancy is linked to behavioural traits (and other non-imaging variables). For this, we applied bayesian partial least square (BPLS) combined with permutation testing, to predict the behavioural variables from states fractional occupancy. The major confounds we controlled for during this analysis are age, sex, head size, and head motion. The confounds are regressed out both from the fractional occupancy and behavioural variables and all reported results are nested 10-fold cross-validated.

We divided the behavioural variables into the following categories: Early life factors, lifestyle and environment (exercise and work), physical measures (general), physical measures (bone density and sizes), physical measures (cardiac and blood vessels), hearing test, blood assays, cognitive phenotypes, health and medical history, genetic measures and mental health self-reported variables. Each category is composed of numerous sub-variables. For instance, the mental health self-reported category consists of variables like risk-taking, frequency of depressed moods, and a number of manic episodes, etc. As shown in Figure S8[A], we demonstrated the non-imaging variables that are predicted using the states fractional occupancy values for the UKB dataset.

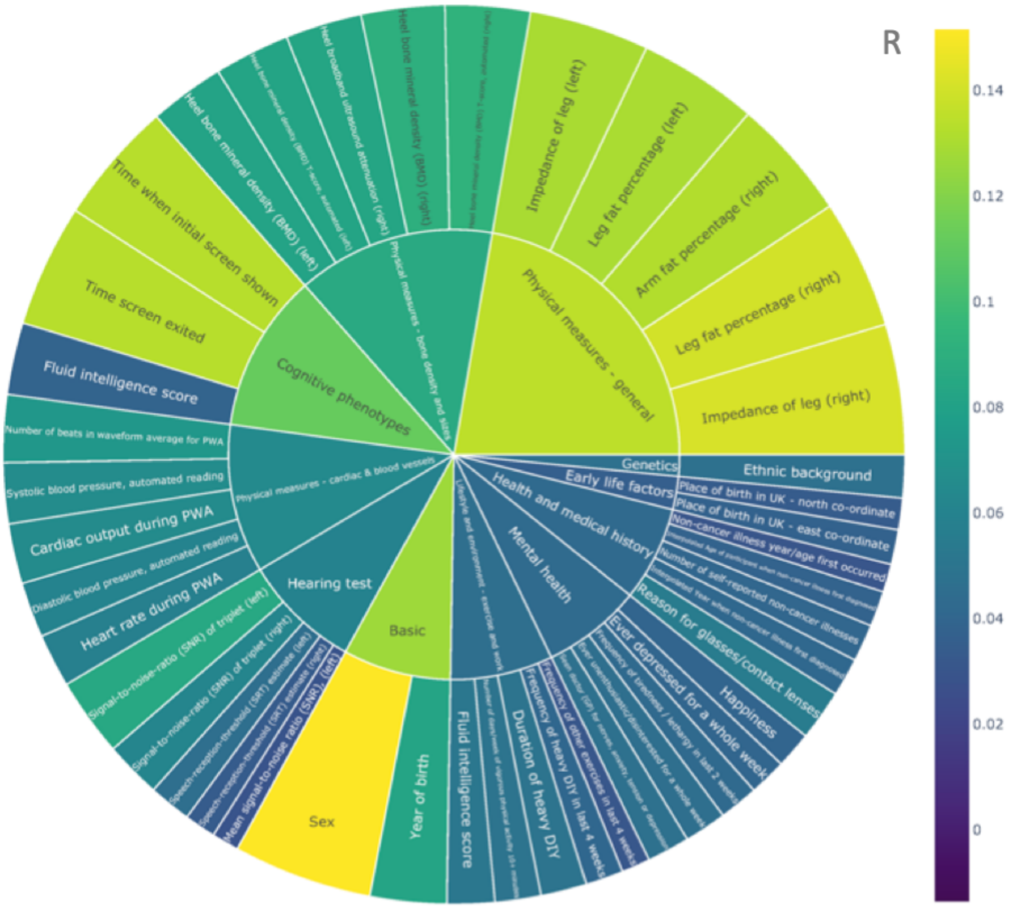


Figure S8: **UKB Dataset:** We have illustrated the amount of correlation explained in the non-imaging behavioural variables using states fractional occupancy. The area of the sub-blocks is colour-coded according to their prediction performance.

#### S2.6. **HCP: MAGE estimated state-specific functional connectivity correlation maps**

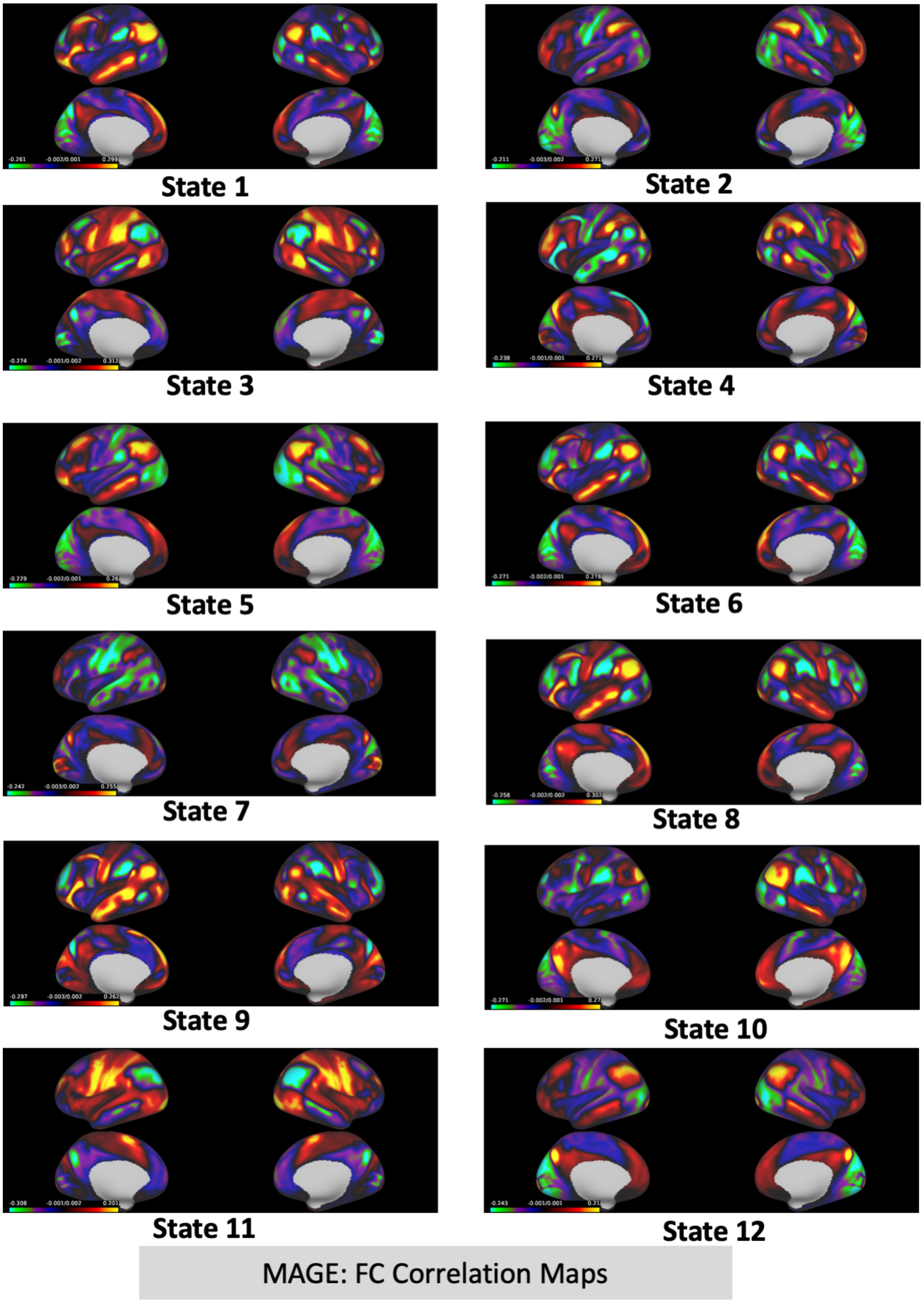


Figure S9: **HCP Data:** We illustrated the spatial maps for the functional connectivity using the multi-dynamic (MAGE) approach for all the 12 states.

#### *S2.7.* ***HCP: MAGE estimated state-specific mean activity maps***

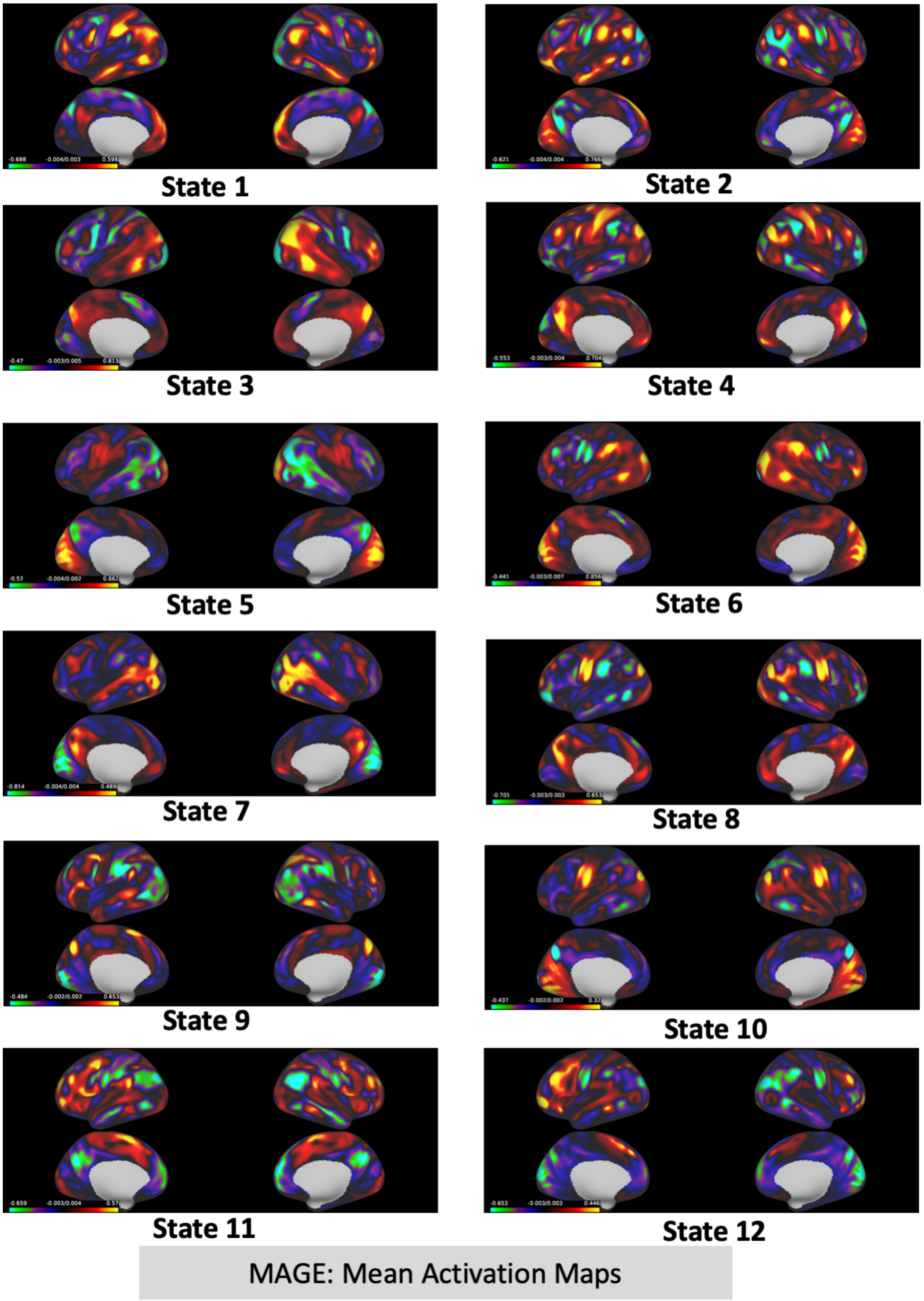


Figure S10: **HCP Data:** We illustrated the spatial maps for the mean activity levels using the multi-dynamic (MAGE) approach for all the 12 states.

#### *S2.8.* ***HCP: SAGE estimated state-specific functional connectivity correlation maps***

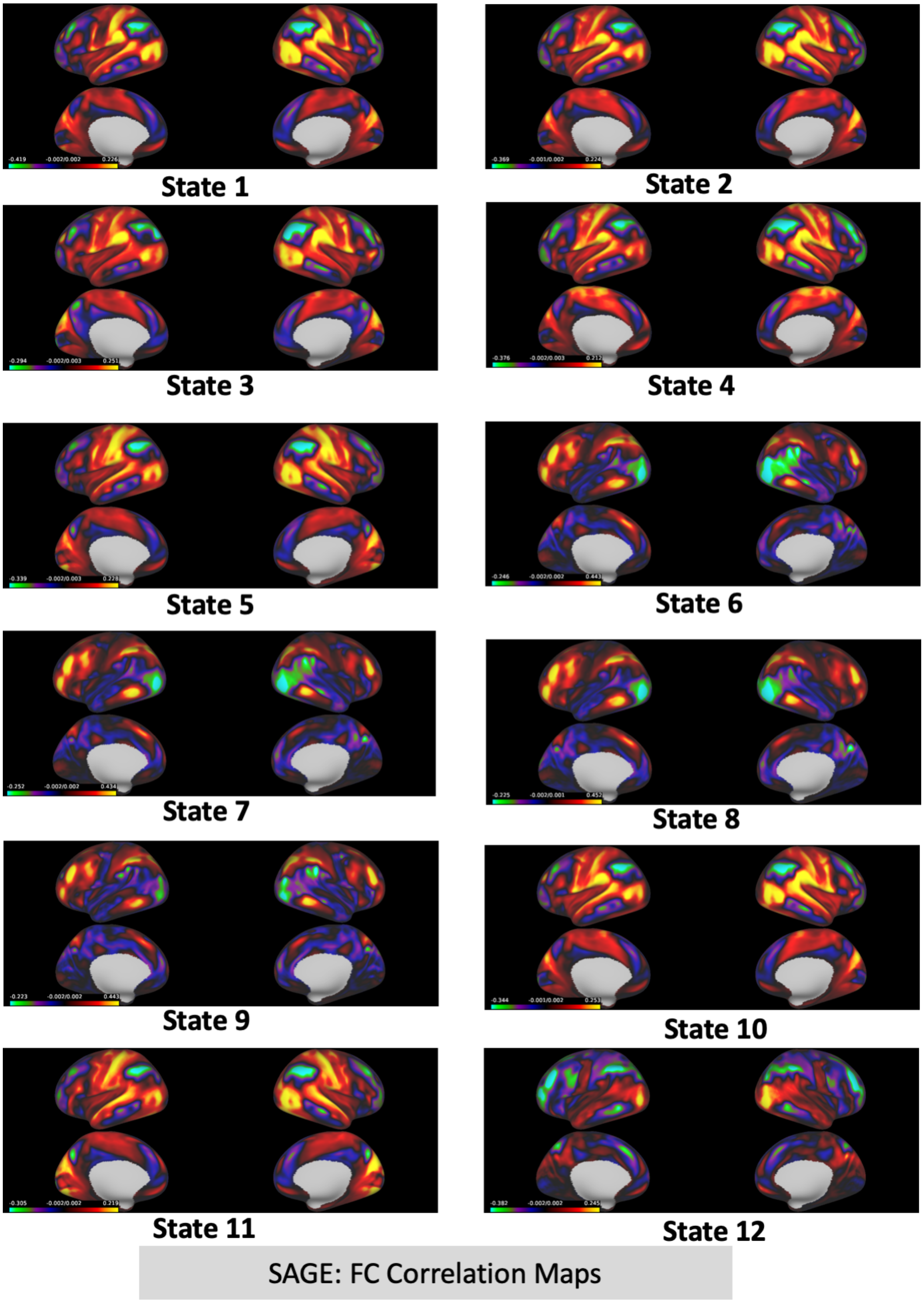


Figure S11: **HCP Data:** We illustrated the spatial maps for the functional connectivity using the single-dynamic (SAGE) approach for all the 12 states.

#### S2.9. ***HCP: SAGE estimated state-specific mean activity maps***

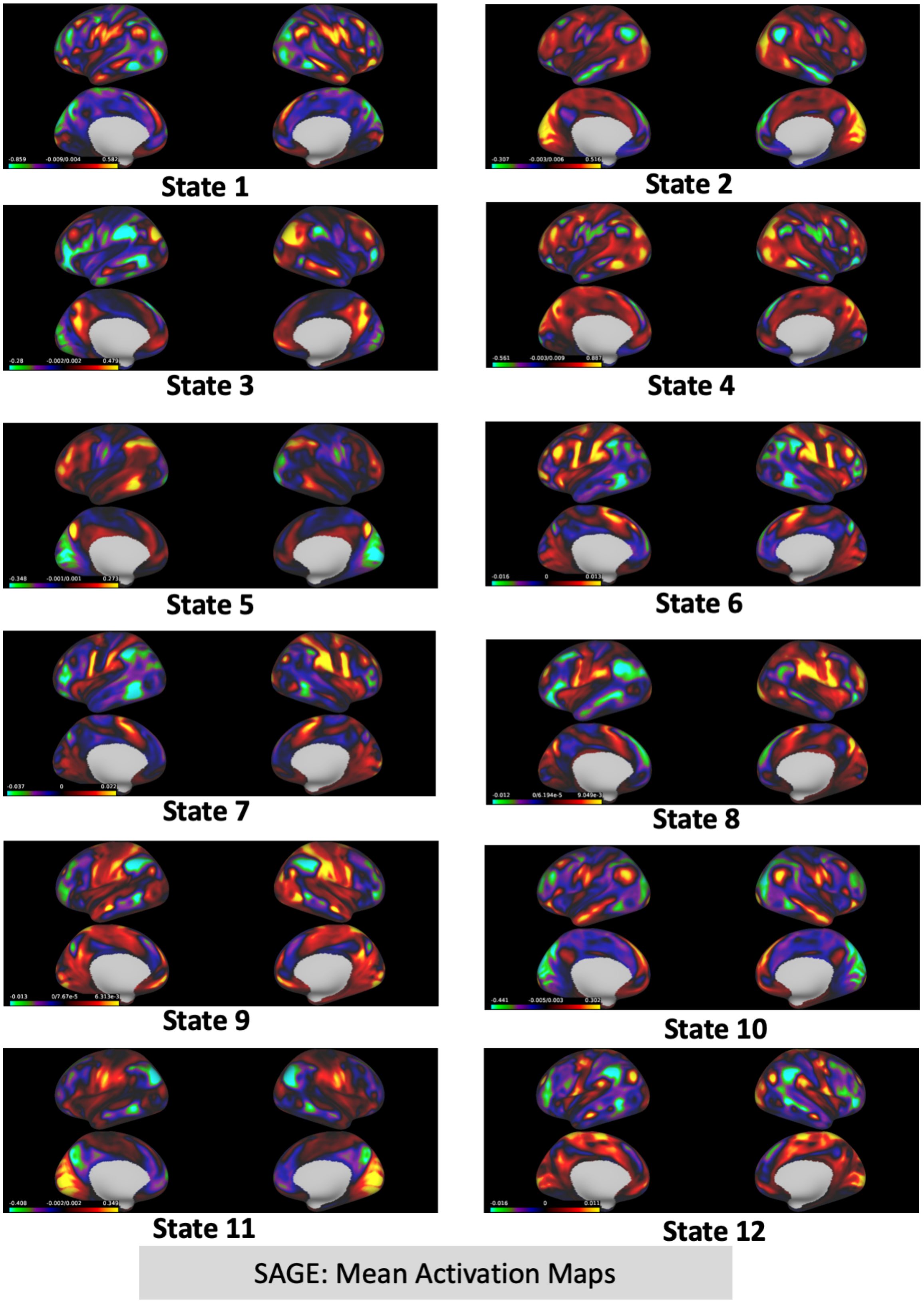


Figure S12: **HCP Data:** We illustrated the spatial maps for the mean activity levels using the single-dynamic (SAGE) approach for all the 12 states.

#### *S2.10.* ***HCP: SAGE estimated state-specific functional connectivity correlation maps (mean activity is forced to zero)***

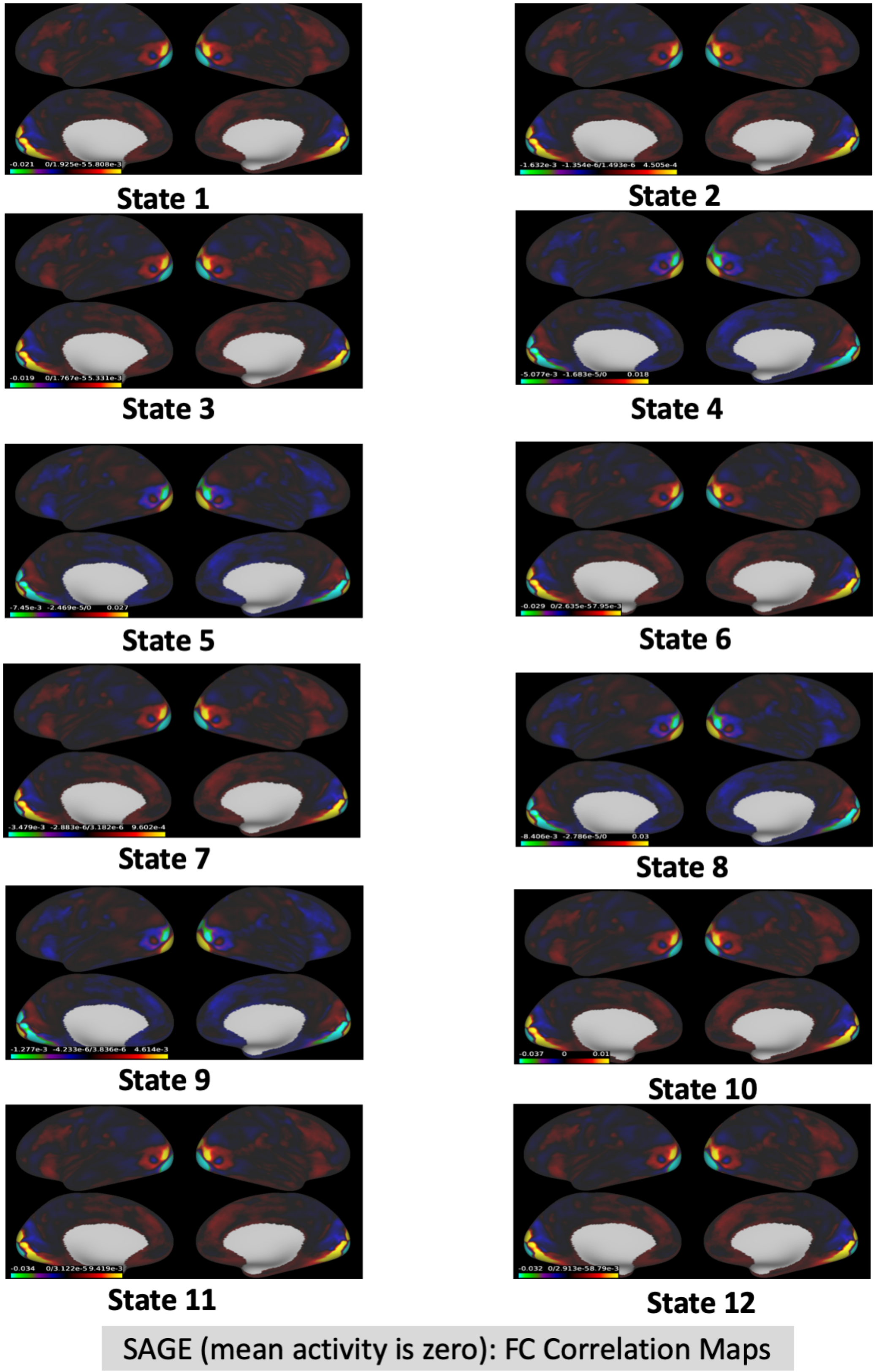


Figure S13: **HCP Data:** We illustrated the spatial maps for the functional connectivity using the single-dynamic (SAGE) approach for all the 12 states.

*S2.11.* ***UKB: SWC estimated state-specific functional connectivity correlation maps***

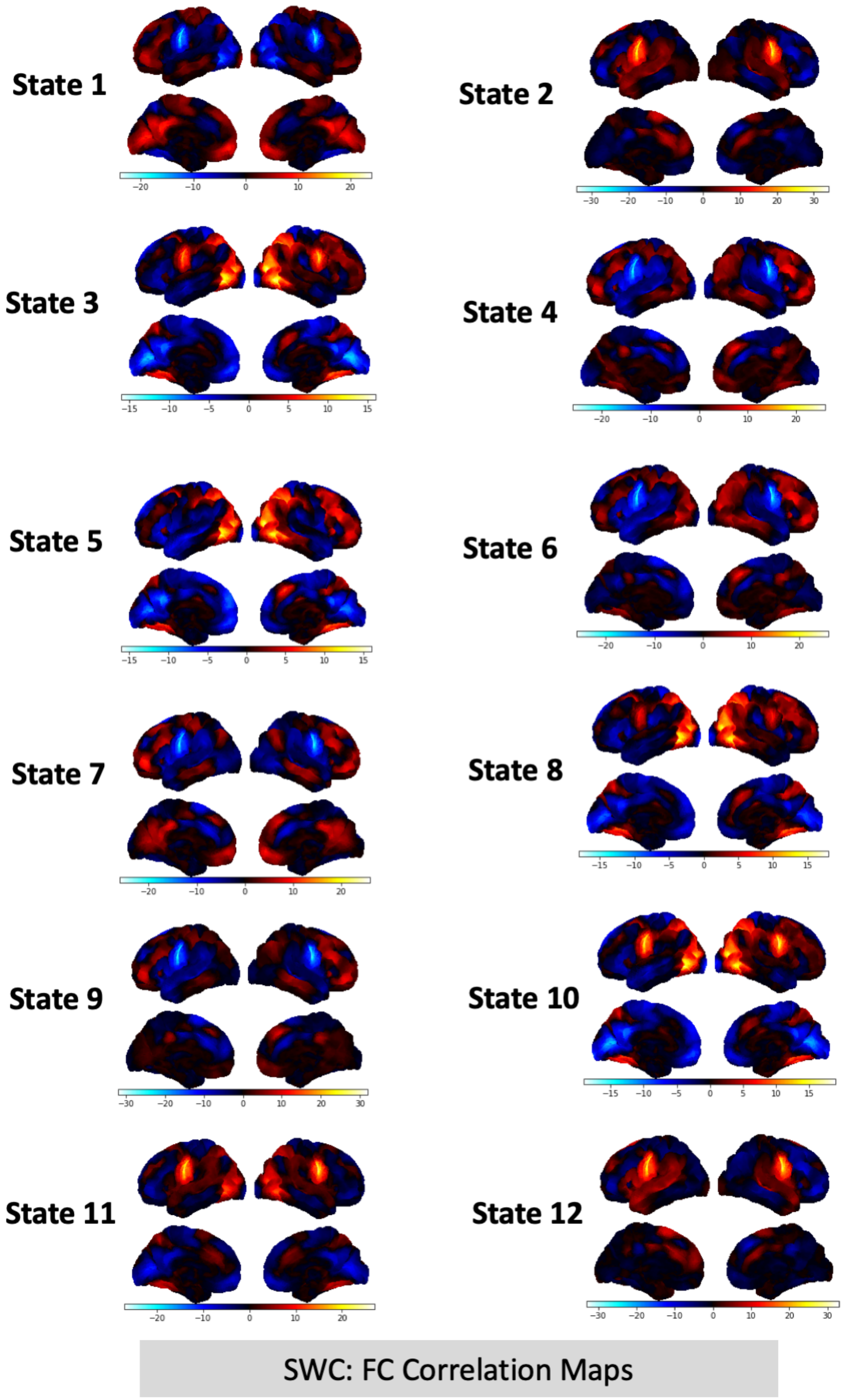


Figure S14: **UKB Data:** We illustrated the spatial maps for the functional connectivity using the Sliding Window Correlation (SWC) approach for all the 12 states.

#### *S2.12.* ***UKB: HMM estimated state-specific functional connectivity correlation maps***

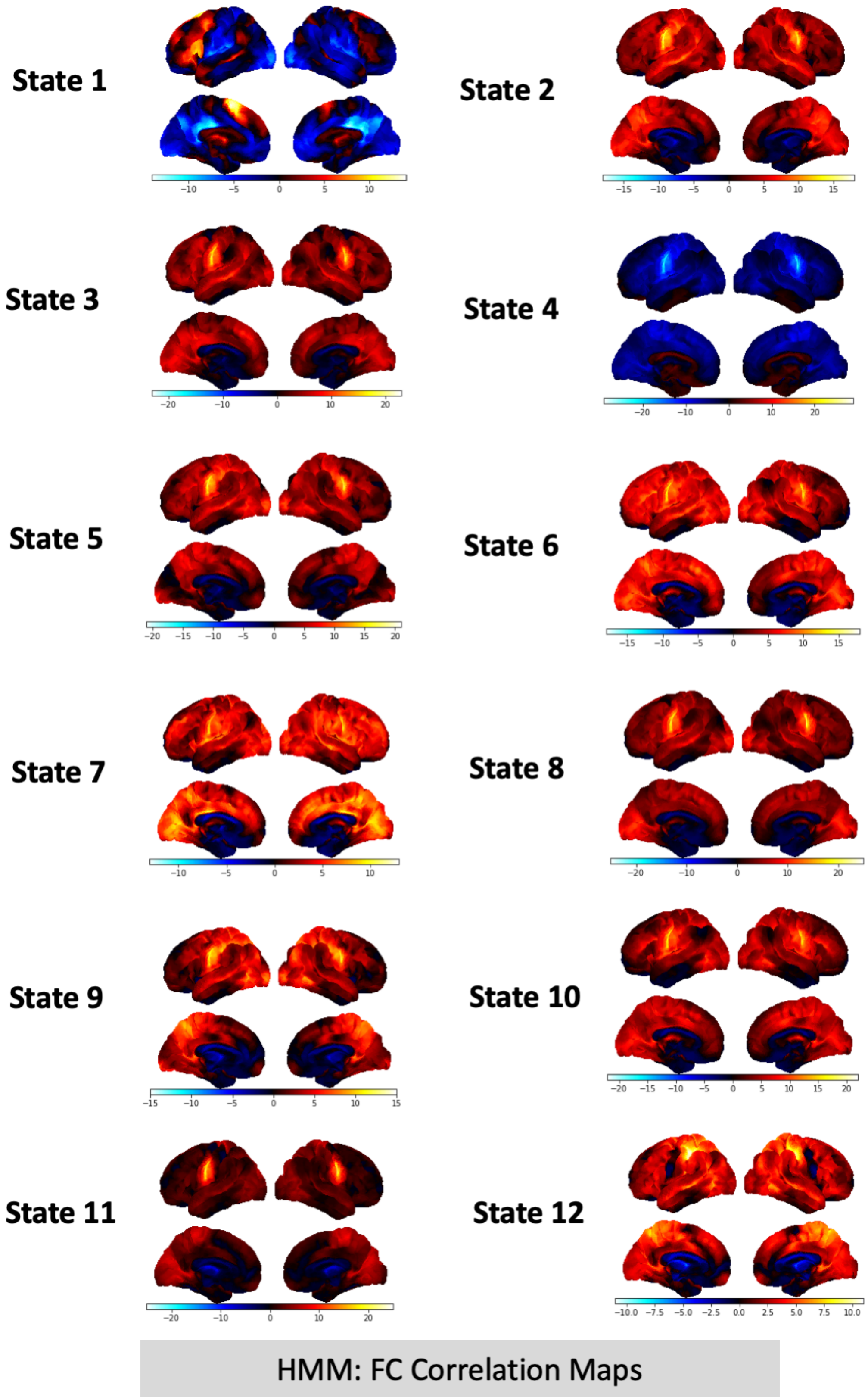


Figure S15: **UKB Data:** We illustrated the spatial maps for the functional connectivity using the Hidden Markov Models (HMM) approach for all the 12 states.

#### *S2.13.* ***UKB: MAGE estimated state-specific functional connectivity correlation maps***

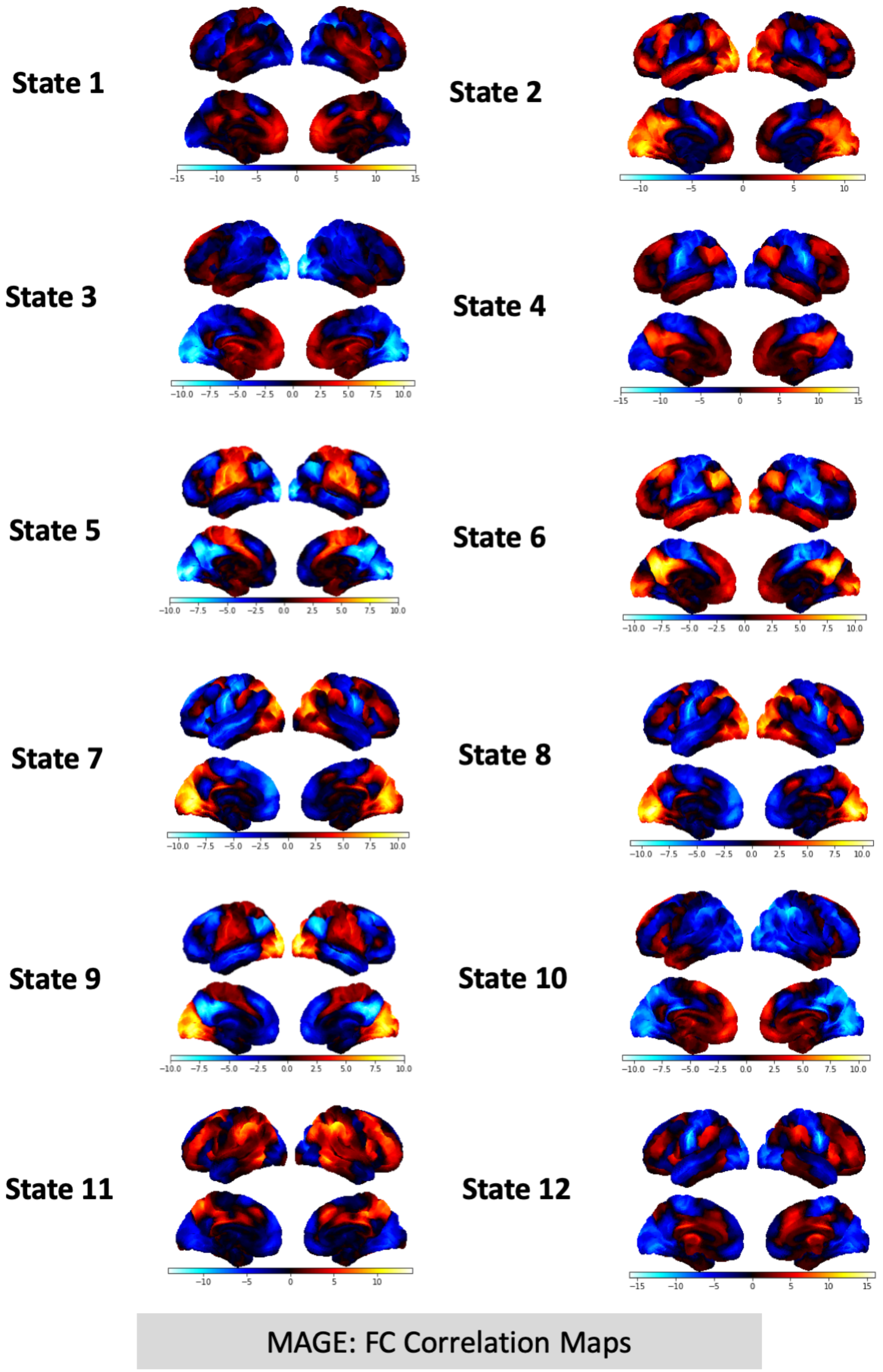


Figure S16: **UKB Data:** We illustrated the spatial maps for the functional connectivity using the multi-time-scale (MAGE) approach for all the 12 states.
